# Human shape representations are not an emergent property of learning to classify objects

**DOI:** 10.1101/2021.12.14.472546

**Authors:** Gaurav Malhotra, Marin Dujmović, John Hummel, Jeffrey S Bowers

**Affiliations:** School of Psychological Sciences, University of Bristol, Bristol, UK; Department of Psychology, University of Illinois Urbana-Champaign, Champaign, USA

## Abstract

Humans are particularly sensitive to changes in the relationships between parts of objects. It remains unclear why this is. One hypothesis is that relational features are highly diagnostic of object categories and emerge as a result of learning to classify objects. We tested this by analysing the internal representations of supervised convolutional neural networks (CNNs) trained to classify large sets of objects. We found that CNNs do not show the same sensitivity to relational changes as previously observed for human participants. Furthermore, when we precisely controlled the deformations to objects, human behaviour was best predicted by the amount of relational changes while CNNs were equally sensitive to all changes. Even changing the statistics of the learning environment by making relations uniquely diagnostic did not make networks more sensitive to relations in general. Our results show that learning to classify objects is not sufficient for the emergence of human shape representations.

## Introduction

A great deal of research into human vision is driven by the observation that visual perception is biased. For example, we prefer to group objects in a scene based on certain Gestalt principles – a bias to look for proximity, similarity, closure and continuity (Ellis, 2013). We also prefer to view objects from certain viewpoints – a bias for canonical-perspectives (Palmer, 1981). This paper is focused on one such bias – the *shape-bias* – the observation that humans, from a very young age, prefer to categorise objects based on their shape, rather than other prominent features such as colour, size or texture (Biederman & Ju, 1988; Landau, Smith, & Jones, 1988). One manifestation of this bias is that we can identify most objects from line drawings as quickly and accurately as we can identify them from full-color photographs (Biederman & Ju, 1988) and we can do this even if we have no previous experience with line drawings (Hochberg & Brooks, 1962).

Two different explanations have been proposed regarding the origin of these biases. The first view, which we call the *heuristic approach*, proposes that biases originate because the visual system needs to transform the *proximal stimulus* – i.e., the retinal image – into a representation of the *distal stimulus* – i.e., a veridical representation of the cause of the stimulus. Of course, the simple act of transforming one representation to another should not necessarily lead to biases. But, in this case, mapping the retinal image to the distal stimulus is an ill-posed problem: there is not enough information in the proximal stimulus to unambiguously recover the properties of the distal stimulus (Nakayama, He, & Shimojo, 1995; Pizlo, 2001). To overcome this problem, the visual system makes assumptions (i.e., employs heuristics) to determine which properties of the proximal stimulus are used to build distal representations (Knill, 1992; Mamassian & Landy, 1998; Pizlo & Stevenson, 1999; Stevens, 1981). A striking example of such assumptions is the Kanizsa triangle (Kanizsa, 1979), where the visual system encodes the multiple collinearities of edges present in the proximal image and uses these to build contours of a triangle even though these contours do not exist in the retinal image. The advantage of distal representations is that they are relevant for a broad range of tasks – the same representation of an object can be used for recognition and visual reasoning (Hummel & Biederman, 1992) amongst other visual skills.

A second view proposes that these biases can emerge as a result of internalisation of the biases present in the environment relevant for classifying objects. According to this view, humans prefer to view objects from a canonical perspective because these perspectives are more frequent in the visual environment, and they prefer to classify objects based on shape because shape is more diagnostic during object classification. In other words, biases are a consequence of performing statistical learning on a large set of objects, with the goal of optimising behaviour on a particular task. We will call this the optimisation-for-classification approach or, more briefly, the *optimisation approach*.

The goal of this study was to test the second view – whether inferences about distal stimuli can emerge as a result of learning to classify a large set of objects. We tested this by focusing on supervised Convolutional Neural Networks (CNNs) – which are machine learning models that recognise objects by learning statistical features of their proximal stimuli that can be used to optimally classify each stimulus, given some training data. The learned representations that support object recognition are specialized for image classification. There is no pressure to learn distal representations of objects. As such, CNNs trained using supervised learning to classify objects provide a concrete model to test the optimisation view. If human perceptual biases are acquired purely through internalising the statistics of the environment in order to classify objects, then training CNNs to perform classification on ecologically realistic datasets should lead to perceptual shape biases similar to the ones observed for humans.

Initial studies testing shape-bias in CNNs showed that CNNs trained in a supervised setting on large datasets of naturalistic images (e.g. `ImageNet`) frequently lacked a shape-bias, instead preferring to classify images based on texture (Geirhos et al., 2018) or other local features (Baker, Lu, Erlikhman, & Kellman, 2018; Malhotra, Dujmovic, & Bowers, 2021). However, it has been argued that CNNs can also be trained to infer an object’s shape given the right type of training. For example, Geirhos et al. (2018) trained standard CNNs on Style-Transfer image dataset that mixes the shape of images from one class with the texture from other classes so that only shape was diagnostic of category. CNNs trained on this dataset learned to classify objects by shape. In another study, Feinman and Lake (2018) found CNNs were capable of learning a shape-bias based on a small set of images, as long as the training data was carefully controlled. Similarly, Hermann, Chen, and Kornblith (2020) showed that more psychologically plausible forms of data augmentation, namely the introduction of color distortion, noise, and blur to input images, make standard CNNs rely more on shape when classifying images. Indeed, the authors found that data augmentation was more effective in inducing a shape bias than modifying the learning algorithms or architectures of networks, and concluded: “Our results indicate that apparent differences in the way humans and ImageNet-trained CNNs process images may arise not primarily from differences in their internal workings, but from differences in the data that they see” (Hermann et al., 2020, Abstract).

These results raise the possibility that human biases are indeed a consequence of internalising the statistical properties of the environment relevant to classifying objects rather than the product of heuristics involved in building distal representations of objects. But studies so far have focused on judging whether or not CNNs are able to develop a shape-bias, rather than examining the type of shape representations they acquire. If humans and CNNs indeed acquire a shape-bias through a similar process of statistical optimisation, then CNNs should not only show a shape-bias, but also develop shape representations that are similar to human shape representations.

A key finding about human shape representations is that humans do not give equal weight to all shape-related features. For example, it has been shown that human participants are more sensitive to distortions of shape that change relations between parts of objects than distortions that preserve these relations (Biederman, 1987; Hummel & Stankiewicz, 1996). These observations have typically been taken to support a heuristic view according to which relations present in the proximal images are used to build distal representations of objects (Hummel, 1994). The question we ask is whether CNNs trained to classify objects learn to encode these relational features of shape. If they do, it would suggest that the relational sensitivity of human shape representations can emerge as a consequence of learning to classify large sets of objects and that shape-biases in object recognition are the product of optimising performance on object classification. But if not, it would suggest that these biases are best characterized as heuristics designed to build distal representations of shape and that learning to classify objects is not sufficient for the emergence of such distal representations.

In the rest of the paper, we discuss a series of experiments (simulation studies with CNNs as well as behavioural experiments with human participants) which show that the shape representations that emerge as a result of classifying images in CNNs are qualitatively different from human shape representations. In the first two experiments, we examine objects that consist of multiple parts, while the following experiments examine objects that consist of a single part. The deformations required to infer the shape representations of these two types of objects are different, but related. Therefore, we begin each section by describing these deformations and how these deformations are predicted to affect shape representations under the two (optimisation and heuristic) views. We then present results of experiments where humans and CNNs were trained on the same set of shapes and then presented these deformations. In the final section, we discuss how our findings pose a challenge for developing models of human vision.

## Experiment 1

In our first experiment, we asked whether models that learn to optimise their performance by classifying large sets of objects develop a key property of human shape representations – it’s sensitivity to a subset of object deformations. According to the structural description theory (Biederman, 1987), humans represent objects as collections of convex parts in specific categorical spatial relations. For example, consider two objects – a bucket and a mug – both of which consist of the same parts: a curved cylinder (the handle) and a truncated cone (the body). The encoding of objects through parts and relations between parts makes it possible to support a range of visual skills. For example, it is possible to appreciate the similarity between a mug and a bucket because they both contain the same parts (curved cylinder and truncated code) as well as their differences (the different relations between the object parts). That is, the representational scheme supports visual reasoning. In addition, the parts themselves are coded so that they can be identified from a wide range of viewing conditions (e.g., invariance to scale, translation and viewing angle, as well as robustness to occlusion), allowing objects to be classified from novel poses and under degraded conditions.

Note that the reliance on categorical relations to build up distal representations of multi-part objects is a built-in assumption of the model (one of the model’s heuristics), and it leads to the first hypothesis we test, namely that image deformations that change a categorical relation between an object’s parts should have a larger impact on the object’s representation than metrically-equivalent deformations that leave the categorical relations intact (as might be produced by viewing a given object from different angles). By contrast, any model that relies only on the properties of the proximal stimulus might be expected to treat all metrically-equivalent deformations as equivalent. Such a model may learn that some distortions are more important – i.e., diagnostic – than others in the context of specific objects, but it is unclear why they would show a general tendency to treat relational deformations as different than metric ones since there is no heuristic that assumes that categorical relations between parts is central feature of object shape representations. (Indeed, it may have no explicit encoding of parts at all.) Instead, all deformations are simply changes in the locations of features in the image.

Hummel and Stankiewicz (1996) designed an experiment to test this prediction of structural description theory and compare it to the prediction of view based models (Poggio & Edelman, 1990) of human vision. They created a collection of shapes modeled on Tarr and Pinker’s (1989) simple “objects”. Each object consisted of a collection of lines connected at right angles (Figure 1). Hummel and Stankiewicz then created two deformations of each of these *Basis* object. One deformation, the *relational* deformation (Rel), was identical to the Basis object from which it was created except that one line was moved so that its “above/below” relation to the line to which it was connected changed (from above to below or vice-versa). This deformation differed from the Basis object in the coordinates of one part and in the categorical relation of one part to another. The other deformation, the *coordinates* deformation (Cood), moved two lines in the Basis object in a way that preserved the categorical spatial relations between all the lines composing the object, but changed the coordinates of two lines. Note that both types of deformations can, in principle, indicate a change in distal stimulus. But, a system that uses relational changes as a heuristic for changes to distal stimuli will be more sensitive to Rel changes than Cood changes.

**Figure 1.**
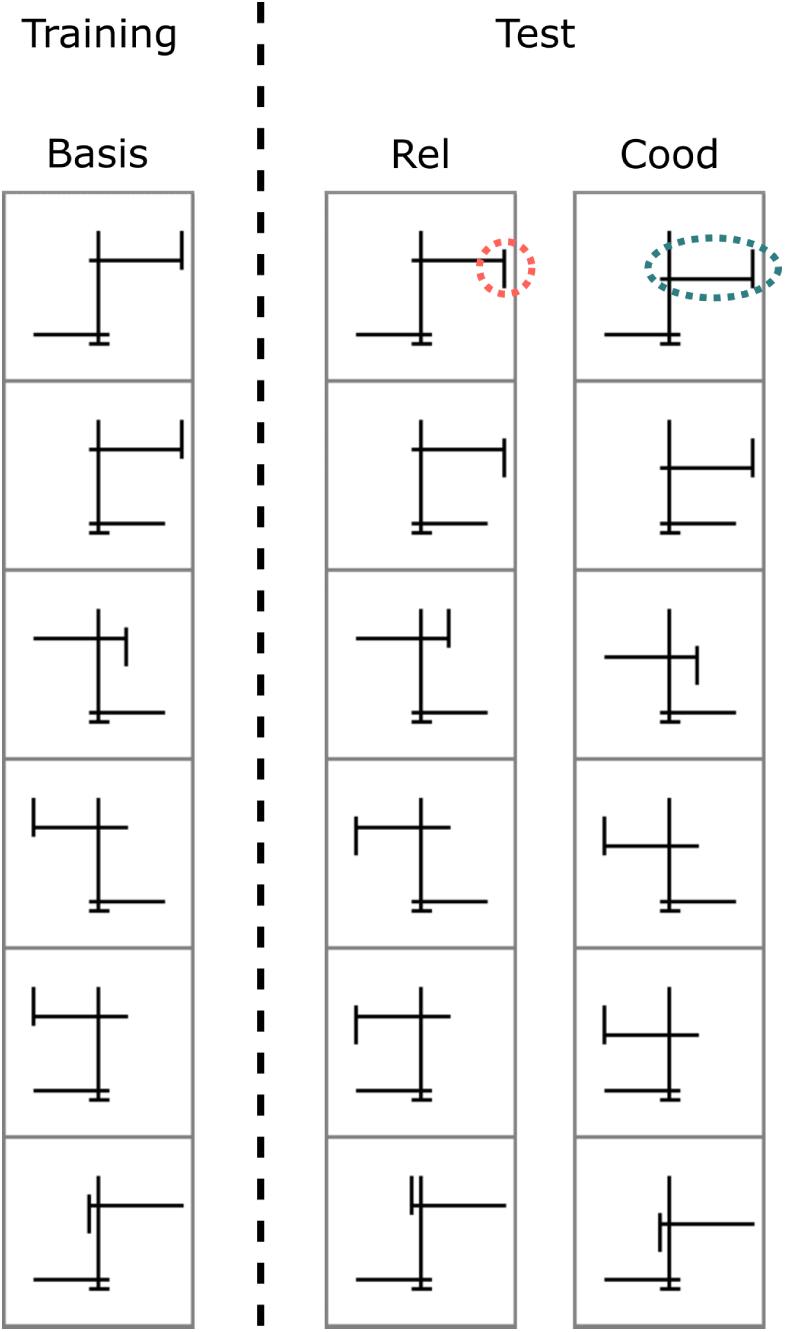
Stimuli used by Hummel and Stankiewicz (1996). *Note.* The first column shows a set of six (Basis) shapes that participants were trained to recognise. Participants were then tested on shapes in the second and third columns, which were generated by deforming the Basis shape in the corresponding row. In the second column (Rel deformation) a shape is generated by changing one categorical relation (highlighted in red circle). In the third column (Cood deformation) all categorical relations are preserved but coordinates of some elements are shifted (highlighted in blue ellipse).

Across five experiments participants first learned to classify a set of base objects and then tested on their ability to distinguish them from their relational (Rel) and coordinate (Cood) deformations. The experiments differed in the specific set of images used, the specific tasks, the duration of the stimuli, but across all experiments, participants found it easy to discriminate the Rel deformations from their corresponding basis object and difficult to distinguish the Cood deformations. The effects were not subtle. In Experiment 1 (that used the stimuli from Figure 1) participants mistook the Rel and Cood images as the base approximately 10% and 90%, respectively, with similar findings observed across experiments. Hummel and Stankiewicz took these findings to support the claim that humans encode objects in terms of the categorical relations between their parts, consistent with the predictions of the structural description theories that propose a heuristic approach to human shape representation (Hummel, 1994).

However, an optimisation approach may also be able to explain the findings of Hummel and Stankiewicz – a bias for perceiving objects in terms of parts and relations may simply *emerge* as a result of learning to classify objects. In Experiment 1, we tested this hypothesis by replicating the experimental setup of Hummel and Stankiewicz, replacing human participants with two well-known CNNs – `VGG-16` and `AlexNet` – that have been previously argued to capture human-like representations (Kriegeskorte, 2015; Yamins & DiCarlo, 2016) and an ability to develop a shape-bias (Geirhos et al., 2018; Hermann et al., 2020).

### Methods

#### Training Stimuli

We constructed six basis shapes that were identical to the shapes used by Hummel and Stankiewicz (1996) in their Experiments 1–3. Each image was sized 196×196 pixels and consisted of five black line segments on a white background organised into different shapes. All images had one short (horizontal) segment at the bottom and one long (vertical) segment in the middle. This left three segments, two long, which were always horizontal, and one short, which was always vertical. The two horizontal segments could be either left-of or right-of the central vertical segment. Additionally, the short vertical segment could be attached to the left-of or the right-of the upper horizontal segment. This means that there were a total of 8 (2×2×2) possible Basis shapes. We selected six out of these to match the six shapes used by Hummel and Stankiewicz (1996). Each training set contained 5000 images in each category constructed using data augmentation, where the Basis image was translated to a random location (in the range [*−*50, +50] pixels) on the canvas and randomly scaled ([0.5, 1]) or rotated ([*−*20°, +20°]).

#### Test Stimuli

Following Hummel and Stankiewicz (1996), we constructed Rel (relational) deformations (called V1 variants by Hummel and Stankiewicz (1996)) of each Basis shape by shifting the location of the top vertical segment, so that it’s categorical relation to the upper horizontal segment changed from “above” to “below”. Similarly, we constructed Cood (coordinate) deformations (called V2 variants by Hummel and Stankiewicz (1996)) by shifting the location of *both* the top horizontal line and the short vertical segments together, so that the categorical relations between all the segments remained the same but the pixel distance (e.g. cosine distance) was at least as large as the pixel distance for the corresponding Rel deformation. The test set consisted of 1000 triplets of Basis, Rel and Cood images for each category, which were again generated using the same data augmentation method.

#### Model architecture and pre-training

We evaluated two deep convolutional neural networks, `VGG-16` (Simonyan & Zisserman, 2014) and `AlexNet` (Krizhevsky, Sutskever, & Hinton, 2012) on the image classification tasks described in the Results section. We obtained qualitatively similar results for both architectures. Therefore, we focus on the results of `VGG-16` in the main text and describe the results of `AlexNet` in Appendix C. Since human participants had a lifetime experience of classifying naturalistic objects prior to the experiment, we used network implementations that had been pre-trained on a set of naturalistic images. Two types of pre-training were used: networks were either pre-trained in the standard manner on `ImageNet` (a large database of naturalistic images), or pre-trained on a set of images where shape was made more predictive than texture by using style-transfer (Gatys, Ecker, & Bethge, 2016). We used networks pre-trained by Geirhos et al. (2018), who have shown that networks trained in this manner have a greater shape-bias than networks trained on `ImageNet`.

#### Further training

Networks were either tested in a *Zero-shot* condition, where no further training was given on any of our datasets and we recorded the response of the pre-trained networks to the test images, or in a *Fine-tuned* condition, where the pre-trained network was fine-tuned to classify the 5000 Basis images of each category described in the Stimuli above. This fine-tuning was performed in the standard manner (Yosinski, Clune, Bengio, & Lipson, 2014) by replacing the last layer of the classifier to reflect the number of target classes in each dataset. The models learnt to minimise the cross-entropy error by using the Adam optimiser (Kingma & Ba, 2014) with a small learning rate of 10*^−^*^5^ and a weight-decay of 10*^−^*^3^. In all simulations, learning continued until the loss function had converged. To check for overfitting, we created cross-validation sets and ensured performance on training set was not higher than on the cross-validation sets. We also trained networks using standard regularization methods such as batch normalization and dropout and obtained qualitatively similar results. In most cases, the networks achieved nearly perfect classification on the training set. All simulations were performed using the `Pytorch` framework (Paszke et al., 2017) and we used `torchvision` implementation of all models.

#### Analysis of internal representations

To test the similarity of internal representations of Basis images and their Rel and Cood deformations, we obtained the embedding of each image at each convolution and fully connected layer of the CNN. For a given category, we randomly sampled 100 pairs of images from the Basis and Rel test sets and computed the cosine similarity between embeddings of each pair. This gave us the estimated average distance in the Ba–Rel condition. Similarly the average cosine similarity between 100 pairs of Basis and Cood test images gave us the Ba–Cood distance. These distances were compared against two baseline conditions. The upper limit of similarity was given by the average similarity of 100 pairs of Basis images from the same category. The lower limit was given by the average similarity of 100 pairs of Basis images from different categories (in each pair, one of the images was from one category and the other from one of the other six categories).

### Results and Discussion

An analysis of the internal representations of `VGG-16` is shown in Figure 2 and it’s classification performance is showin in Figure A1 in Appendix A. (Results for `AlexNet` followed the same qualitative pattern and are shown in Appendix C). Each panel in Figure 2 corresponds to a combination of pre-training and test conditions and shows the average cosine similarity between internal representations for a Basis image and it’s relational (Ba-Rel, solid red line) and coordinate (Ba-Cood, dashed blue line) deformations. The internal representations are computed at all convolutional and fully connected layers within the network. We compared these similarities to two baselines: the average similarity between two Basis images that belong to the same category and the average similarity between two Basis images that belong to different categories. These two baselines provide the upper and lower bounds on similarities (hatched yellow region).

**Figure 2.**
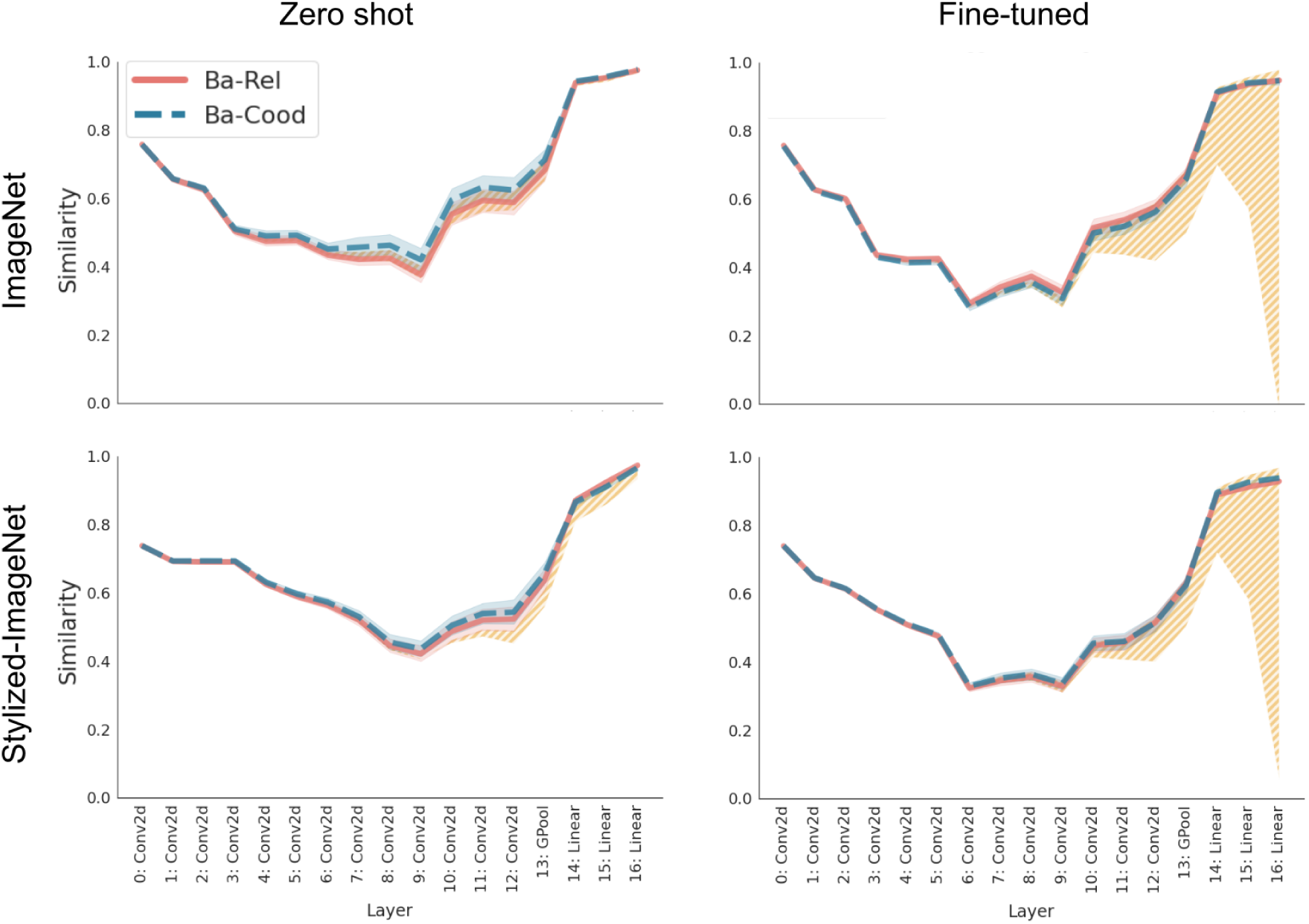
Cosine similarities in internal representations for a VGG-16 network. *Note.* In each panel, the solid (red) line plots the cosine similarity between the internal representations of a Basis shape and its Rel deformation, while the dashed (blue) line plots the cosine similarity between the internal representations of a Basis shape and its Cood deformation. Layers of the network are along the x-axis with Conv2d Linear indicating convolutional and fully connected layers, respectively. Networks were either pre-trained on `ImageNet` (first row) or on `Stylized-ImageNet` – a dataset developed to induce a shape-bias in CNNs (second row). Their internal representations were then probed either without any further training (Zero-shot, first column) or fine-tuning on the Hummel and Stankiewicz dataset (second column). The hatched (yellow) area shows the upper and lower bounds on cosine similarity (obtained by computing the cosine similarity of images from the same and different categories, respectively). Shaded regions around each line show 95% confidence interval. Based on the results of Hummel and Stankiewicz (1996), we would expect the solid (red) line (Ba-Rel) to be closer to the lower, rather than upper bound. Instead we observe that it stays at the upper bound throughout the network and overlaps the dashed (blue) line (Ba-Cood).

We observed that in the *Zero-shot* condition (left-hand column in Figure 2), the similarity between a basis image and it’s relational variant was the same on average (across seeds) as the similarity between the basis image and it’s coordinate variant throughout the networks. That is, the networks failed to distinguish between the basis images and their relational and coordinate variants. In fact, networks also failed to distinguish between basis images from different categories (note the narrow hatched (yellow) region in the *Zero shot* condition in Figure 2 and the classification performance in Figure A1). Thus pre-training on `ImageNet` or `Stylized-ImageNet` was not sufficient for networks to distinguish between the stimuli or their deformations used by Hummel and Stankiewicz – to these models, all line drawings are alike. In contrast, the networks successfully learned to distinguish between stimuli from different categories in the *Fine-tuned* condition (see classification performance in Figure A1). Examining the internal representations showed that the networks represented all types of images in a similar manner in the early convolution layers (there is no difference between similarities within or between categories in the early layers) but representations begin to separate in the deeper convolution and fully connected layers (the hatched (yellow) region increases in size as we move left to right because images from different categories have lower similarity than images from the same category). However, both types of pre-treained networks, the basis images were equally distant to their relational and coordinate deformations (see the overlapping Ba-Rel and Ba-Cood lines in Figure 2). Note that this is not because the networks overfit to the training data. In fact, networks showed very good generalisation to both novel Basis images (in unseen combinations of rotations, translation and scale) and the two types of deformations, with the cosine distance between a basis shape and either deformation close to the upper bound of similarity (also see the high classification performance for both deformations in Figure A1). In summary, we did not find any evidence that suggests that the CNN represents a relational change to an image in any privileged manner compared to a coordinate change.

## Experiment 2

The results of Experiment 1 suggested that learning to classify the naturalistic images in `ImageNet` or even `Stylized-ImageNet` is not sufficient for CNNs to perceive the objects in terms of their categorical relations. But it could be argued that this is not because of a limitation of the optimisation approach, but due to the limitation of datasets that the model was trained on. It is possible that, if the classification model was trained on datasets where relational differences were diagnostic of object categories, it may have internalised this statistic and started perceiving objects in terms of their categorical relations, just like humans. We tested this hypothesis in the next set of simulations, where we created a training environment with a “relational bias”. We show next that when we do this, the network can learn specific changes to relations but it does not generalise this knowledge to novel (but highly similar) relational changes.

### Methods

Experiment 2 used the same model architectures, pre-training and analysis methods as Experiment 1. However, instead of using the training dataset based on Hummel and Stankiewicz (1996), we created three new datasets where relational changes were diagnostic of image categories.

#### Training and Test Stimuli

We generated three datasets – shown as Set 1, Set 2 and Set 3 in Figure 3 – for teaching CNNs to recognise relational deformations on Hummel and Stankiewicz’s stimuli. Each dataset again contained the six Basis shapes (and their translation, rotation and scale transformations) from the training set in Experiment 1. Additionally, they also contained five new Basis shapes. These five shapes were the relational (Rel) deformations of the first five Basis shapes. In other words, the training set assigned different categories to a shape and it’s Rel deformation for five out of six figures. The test set consisted of Rel and Cood deformations of the final (unpaired) shape. Each training set again consisted of 5000 images per category, where each image was constructed by translating, scaling and rotating the Basis shape for that category. The test set consisted of 1000 images per category where each image was constructed by randomly translating, scaling and rotating the Rel and Cood deformations of the unpaired Basis shape.

**Figure 3.**
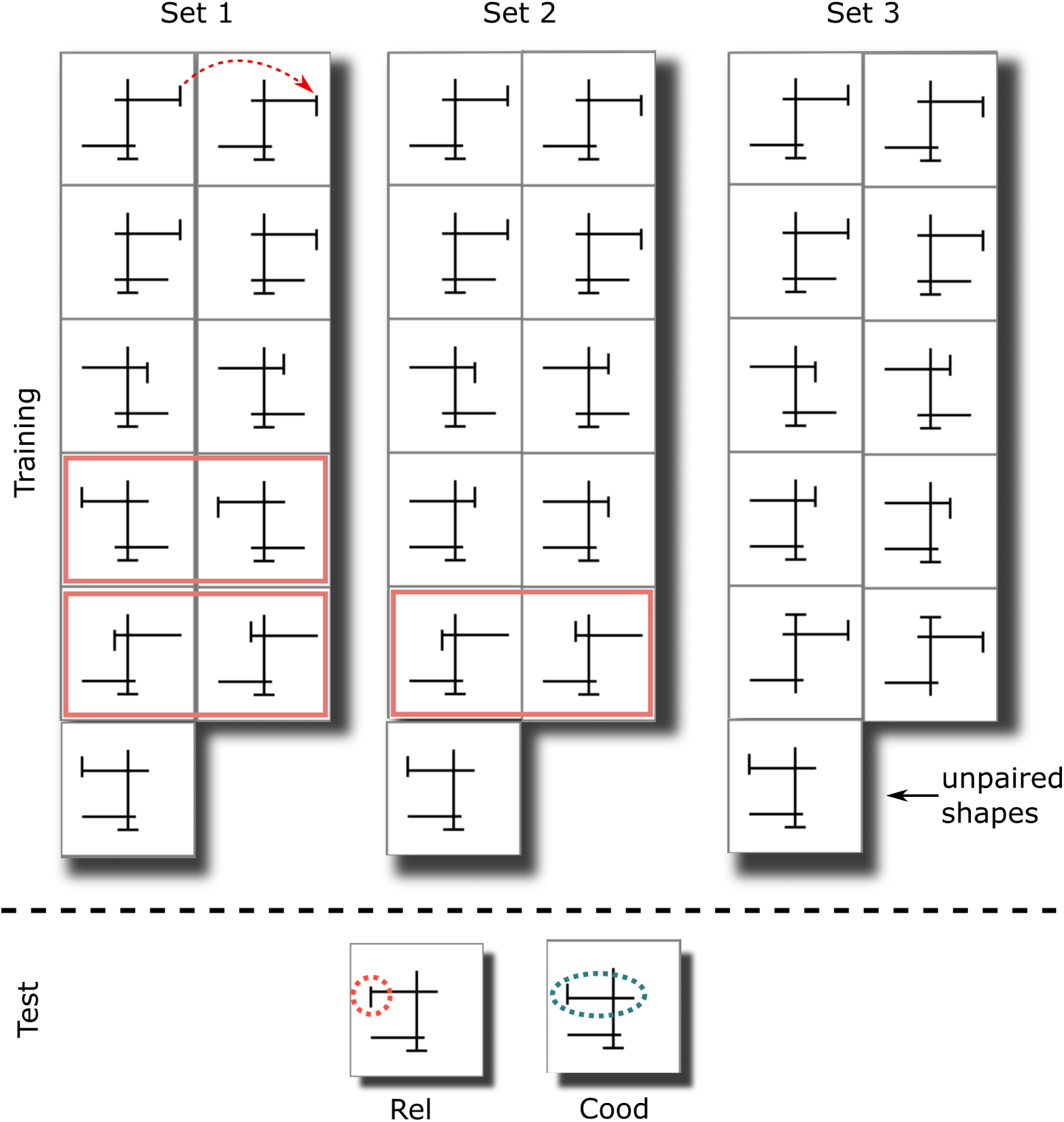
Three training sets that try to teach the network to recognise relational changes. *Note.* In each set, the first column shows a set of six unique Basis shapes, while the second column shows Rel deformations of the first five shapes (see red arrow). At the bottom are the two test shapes. These test shapes are identical to the eleventh (unpaired) training shape, except for one relational (dashed red circle) or coordinate (dashed blue ellipse) deformation. In Set 1, the difference between the untrained shape and the tested Rel deformation is exactly the same as the relational change distinguishing one pair of shapes and similar to another pair in the training set (both highlighted in solid red rectangles). In Set 2, the exact relational change is not trained, however there is similar relational change at a close location (pair again highlighted in solid red rectangle). Set 3 is the most challenging, where none of the diagnostic relational changes in the training set occur at similar locations to the tested relational deformation.

The difference between Set 1, 2, and 3 lay in the degree of novelty of test images. In all three datasets the same relation (dashed red circle in Figure 3) was changed between the unpaired Basis shape and it’s Rel deformation. However, in the first set, there were four other categories (two pairs, highlighted in red rectangles) in the training set where a similar change in relation occurred – that is, for all highlighted categories, there existed another category where the short red segment at the left end of the top bar flipped from “above” to “below” or vice-versa. In the second training set (Set 2 in Figure 3) there were two categories in the training set where the tested relation changed. However, in this case, this relational change occurred in a different location (closer to central vertical line). In the third training set (Set 3 in Figure 3), the tested relational change was the least similar to training (relational changes only occurred to the right of central vertical line for all other trained images).

#### Further Training

Like in Experiment 1, the CNNs were fine-tuned on the training stimuli consisting of eleven (5 + 5 + 1) Basis shapes. All other details of training, including hyper-parameters were kept the same as in Experiment 1.

### Results and Discussion

Figure 4 shows the cosine similarity in internal representations for `VGG-16` trained on these three modified data sets (we obtained a similar pattern of results for `AlexNet` – see Figure C3). As in previous simulations, we tested networks that were either pre-trained on `ImageNet` (first row) or on `Stylized-ImageNet` (second row) and fine-tuned to each training set. We observed that when networks were trained on Set 1 (left column in Figure 4), the cosine similarity Ba-Rel was lower than Ba-Cood in deeper layers of the CNN. That is, the networks treated the relational deformation as *less* similar to Basis figures than the coordinate deformations. This looks much more like the behaviour of human participants in Hummel and Stankiewicz (1996). But note that Set 1 contained two pairs of categories with the same relational change that distinguishes the tested Rel deformation from the corresponding Basis figure. A stronger test is provided by Set 2 that excludes the pair of categories distinguished by the critical relational change from the training set. Here, we observed that the this effect was significantly reduced (middle column in Figure 4) – the cosine similarity Ba-Rel was slightly lower than Ba-Cood but by a much smaller degree and the difference existed only for the networks pre-trained on `ImageNet` and only in the fully connected layers (also compare results in Figure C3 in Appendix for `AlexNet`, where this effect is slightly more pronounced but qualitatively similar). The strongest test for whether the network learns relational representations is provided by Set 3, where none of the categories in the training set changed the exact relation that distinguishes the Rel deformation from the Basis image in the test set. Here, we observed (Figure 4, right-hand column) that the effect disappeared completely – the cosine similarity Ba-Rel was indistinguishable from Ba-Cood and both similarities were at the upper bound. All networks failed to learn that novel relational changes are more important for classification than coordinate changes even when the learning environment contained a “relational bias” – i.e., changing relations led to a change in an image’s category mapping.

**Figure 4.**
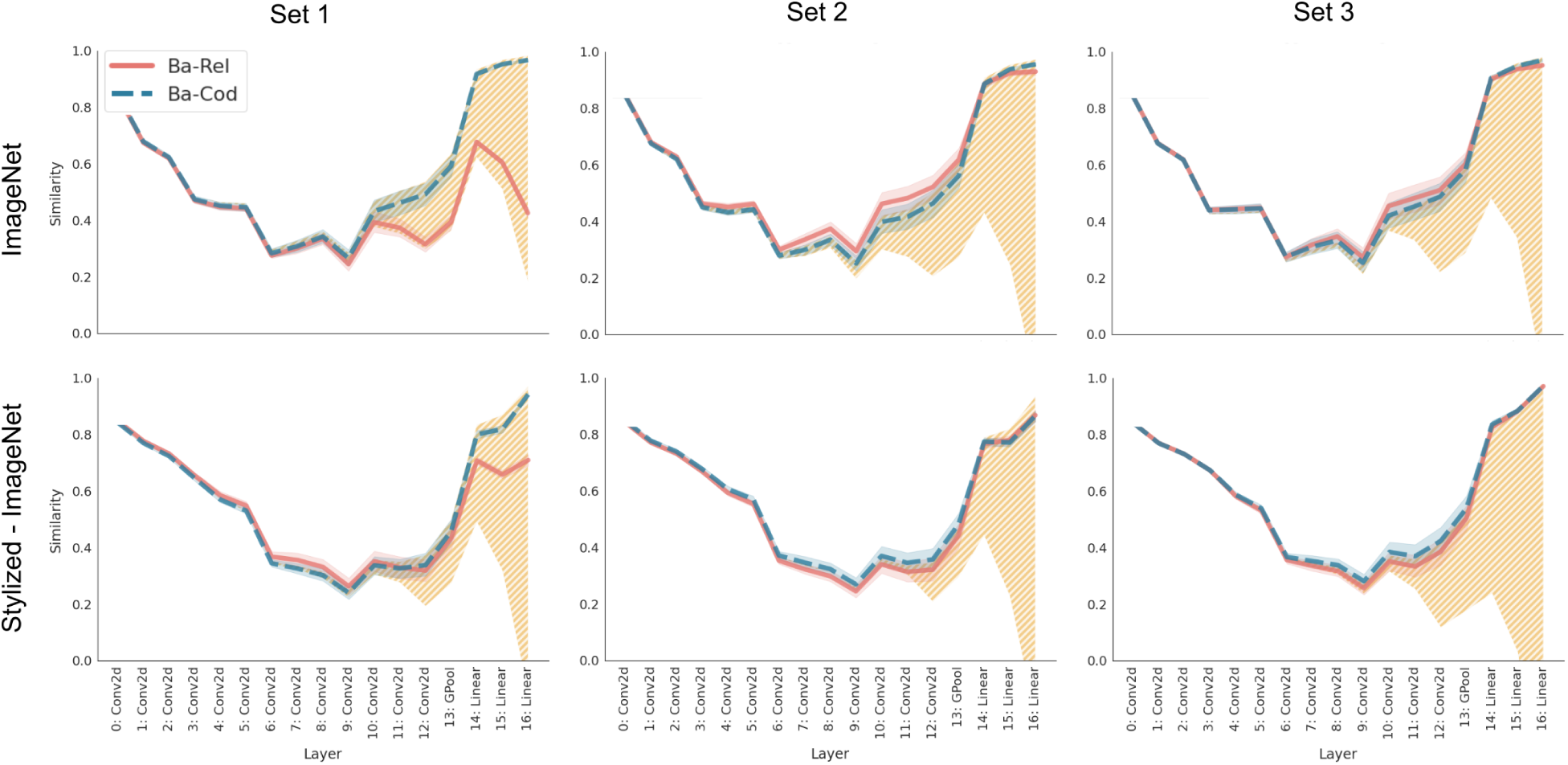
Cosine similarity for VGG-16 networks that have been trained on diagnostic relations. *Note.* Each panel shows cosine similarity in internal representations between Basis images and Rel (solid, red) or Cood (dashed, blue) deformations of those images. The network was either pre-trained on `ImageNet` (first row) or `Stylized-ImageNet` (second row) and fine-tuned on Set 1 (left), Set 2 (middle) or Set 3 (right) shown in Figure 3. Like Figure 2, the hatched (yellow) region shows the upper and lower bound on similarity. We can see that the network fine-tuned on Set 1 represents relational deformations as significantly different from Basis images as well as coordinate deformations (solid red line is much lower than upper bound and dashed blue line for deeper layers in the network). However, this is not the case for networks fine-tuned on Set 2 or Set 3.

## Experiment 3

Experiments 1 and 2 used stimuli that consisted of multiple parts and relational deformations involved changing the relationships between these parts. But of course, in order to build distal representations of complex objects, it is also necessary to build distal representations of the parts themselves. While Experiments 1 and 2 show that CNNs and humans differ in their representation of multi-part objects, the stimuli used in these experiments did not allow us to compare the representations of the parts themselves, or indeed single-part objects. Another limitation of the stimuli in Experiments 1 and 2 was that it used discrete relations (‘up’ vs ‘down’, ‘left of’ vs ‘right of’), which do not permit a manipulation of the degree of relational or coordinate change. This meant that we could not match the extent of relational change with *an equivalent* coordinate change and compare the sensitivity to each of these changes.

What sorts of deformations of the proximal stimulus should allow us to contrast optimisation and heuristic approaches for identifying the component parts of complex objects or single-part objects? According to the structural description theory (Biederman, 1987), certain shape properties of the proximal image are taken by the visual system as strong evidence that individual parts have those properties. For example, if there is a straight or parallel line in the image, the visual system infers that the part contains a straight edge or parallel edges. If the proximal stimulus is symmetrical, it is assumed that the part is symmetrical (see, for example, Pizlo, Sawada, Li, Kropatsch, & Steinman, 2010). These (and other) shape features used to build a distal representation of the object part are called non-accidental because they would only rarely be produced by accidental alignments of viewpoint. The visual system ignores the possibility that a given non-accidental feature in the proximal stimulus (e.g., a straight line) is the product of an accidental alignment of eye and distal stimulus (e.g., a curved edge). That is, the human visual system uses non-accidental proximal features as a heuristic to infer distal representations of object parts. Critical for our purpose, many of the non-accidental features described by Biederman (1987) are relational features, and indeed, many of the features are associated with Gestalt rules of perceptual organization, such as good continuation, symmetry, and Pragnanz (simplicity). Accordingly, any deformations of the proximal stimulus that alter these non-accidental features (such as disrupting symmetry) should have a larger impact on classifications than deformations that do not.

With this in mind, we designed a new stimuli set that allowed us to precisely manipulate the relational and coordinate deformations of single-part objects. The stimuli set consisted of seven symmetrical polygons (see Figure 5(a)), and we deformed these polygons by altering the locations of the vertices composing the polygons in a way that precisely controlled the metric change in the vertices’ locations (in the retinal image). Like Experiment 1, we created two types of deformations: (a) a coordinate deformation that parametrically varied the degree to which a polygon rotated in the visual image, vs. (b) a relational change that had an equivalent impact as the corresponding rotation, but instead introduced a shear that changed relative location of the polygon’s vertices. Note that rotating an object preserves all non-accidental features (Biederman, 1987), while shearing it changes it’s symmetry – a non-accidental property of the object. To a model that looks only at proximal stimulus, both deformations lead to an equivalent pixel-by-pixel change, while to a model that infers properties such as symmetry and solidity of the distal stimulus, the coordinate deformation preserves these properties while the relational deformation changes them.

**Figure 5.**
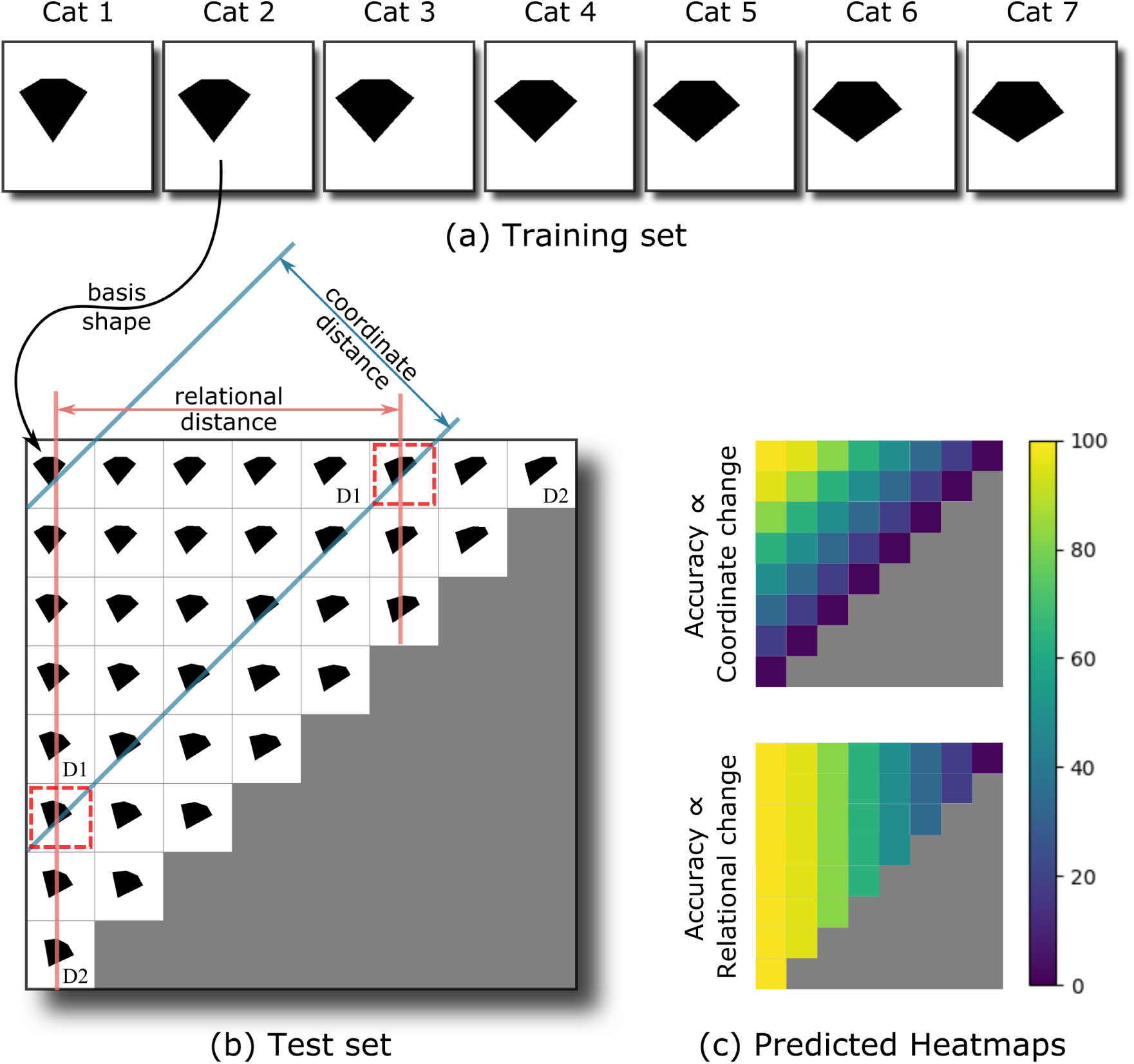
Stimuli used to test shape representations in single-part objects. *Note.* (a) The shapes in the Basis set used for training. Each shape is presented at various translations and scales. (b) The test set for one of the categories (Cat 2) is obtained by deforming the Basis shape (in the top-left corner) through a combination of rotation and shear operations. Here we have organised these deformations in a matrix based on their coordinate distance (measured as cosine distance) and relational distance (measured as change in relative location of vertices) from the basis shape. All deformations on a diagonal of this matrix are at the same coordinate distance from the Basis shape and all deformations in a column are at the same relational distance from the Basis shape. Highlighted (red) squares show stimuli for computing cosine distance in Figure 7 below. Deformations marked D1 and D2 are used for testing human participants. (c) The predicted accuracy on the test set presented as heat-maps, assuming that accuracy is a function of coordinate distance (top), or relational distance (bottom).

Figure 5(b) shows some examples of test images for one of the trained shapes. These test shapes are organised based on the degree and type of deformation. The degree of relational deformation (shear) of a test image increases as we move from left to right, while the degree of coordinate deformation (rotation) increases as we move from top to bottom. We can also construct test shapes that are a combination of these relational and coordinate deformations. Every shape in Figure 5(b) is a combination of a rotation and a shear of the basis shape in the top-left corner. We have organised these test shapes based on their distance to the basis figure: all shapes along each diagonal have the same cosine distance to the basis shape^1^, and diagonals farther from the basis shape are at a larger distance. Thus, this method gives us a set of test shapes organised according to increasing relational and coordinate changes and matched based on the distance to the basis shape. We could now ask how accuracy degrades on this landscape of test shapes. If the visual system encodes shape as a set of diagnostic features of the proximal (retinal) image, accuracy should fall as one moves across (perpendicular to) the diagonals on the landscape. On the other hand, if the visual system encodes shape as a property of the distal stimulus, then changing internal relations should lead to a larger change in classification accuracy than an equivalent coordinate change – that is, the accuracy should fall sharply as one moves left to right along each diagonal. Figure 5(c) shows predicted accuracy on this landscape for the two types of shape representations.

### Methods

#### Training Stimuli

The training set for Experiment 2 consisted of seven symmetric filled pentagons, presented on a white canvas. Each category contained 5000 training images. The training set presented these polygons at different translations and scales, so it was not possible to classify them based on the position of a local feature or the area of the polygon. The difference between Basis shapes for two categories was the angles between the edges. Note that all polygons in the training set were presented in the upright orientation since rotation is the one of the transforms (the coordinate transform) that the model is tested on.

#### Test Stimuli

The test set consisted of a grid of shapes that were obtained by deforming the Basis shape of the corresponding category. We used two deformations: rotation, which preserved the internal angles between edges, and shear, which changed internal angles. To shear a shape, it’s vertices were horizontally moved by a distance that depended on the vertical distance to the apex. For a vertex with coordinates (*x_old_, y_old_*), we obtained a new set of vertices, (*x_new_, y_new_*) = (*x_old_* + *λ*(Δ*y*)^2^*, y_old_*), where *λ* was the degree of shear and Δ*y* was the distance between *y_old_* and *y_apex_*, the y-coordinate of the vertex at the apex. Images could also be combination of rotations and shears. To do this, the Basis image was first sheared, then rotated. We measured the (cosine or Euclidean) distance of a deformed image from the Basis image and used this distance to organise the test images on a grid (see Figure 5), where images in each column had the same degree of shear and images along each diagonal had the same (cosine or Euclidean) distance to the Basis image. We then obtained 100 exemplars of each deformed image on the grid by randomly translating and scaling the image.

#### Model architecture and pre-training

We used the same set of models as Experiments 1 and 2 (`VGG-16` and `AlexNet`) pre-trained in the same manner (either on `ImageNet` or on `Stylized-ImageNet`).

#### Further training

Like Experiments 1 and 2, models were tested in either the *Zero-shot* condition, where we did not train the model on our training set and simply examined the internal representations in response to test images, or in the *Fine-tuned* condition, where the pre-trained model underwent further training (with a reduced learning rate) on the training stimuli. We again observed that the models failed to distinguish any shape in the Zero-shot condition, therefore we restrict the presentation of our results to the Fine-tuned condition.

#### Analysis of internal representations

The similarity of internal reprsentations for the polygons stimuli is obtained in a similar manner to the Hummel and Stankiewicz stimuli. The similarity of a shear transformation to the corresponding Basis image (Ba-Sh) is estimated by measuring the average cosine similarity between embeddings (at all convolutional and fully-connected layers of the network) of 100 pairs images from the Basis and sheared sets of the same category. Similarly, the similarity between a Basis image and it’s rotational transformation (Ba-Rot) is estimated by measuring the average cosine similarity between embeddings of 100 pairs of images from Basis and rotated sets of the same category.

### Results and Discussion

The classification performance of `VGG-16` for images from the test set is shown in Figure 6 (we obtained a qualitatively similar pattern of results for `AlexNet`, see Appendix C). For all networks, we observed that test accuracy was highest at the top-left corner (i.e., for the Basis shape) and reduced as the degree of relational and coordinate change was increased. Thus, unlike Experiment 1, where we were able to observe only ceiling performance for both deformations, the design of Experiment 3 allowed us to compare how performance degrades for the two types of deformations. Crucially, we observed that for most categories, accuracy decreased as a function of distance to the Basis shape (perpendicular to the diagonals), rather than relational change (left to right). In fact, for some categories accuracy *improved* as one moved from left to right along the diagonals. Occasionally, we observed high accuracy for large rotations on one category. This was generally due to false positives, where large rotations for all categories were classified as the same category by the network (see Figure B1 in Appendix B for details). Overall, these results suggest that the network does not represent the shapes in this task in a relational manner. If it did, it’s performance on relational changes should have been a lot worse than it’s performance on relation-preserving rotations.

**Figure 6.**
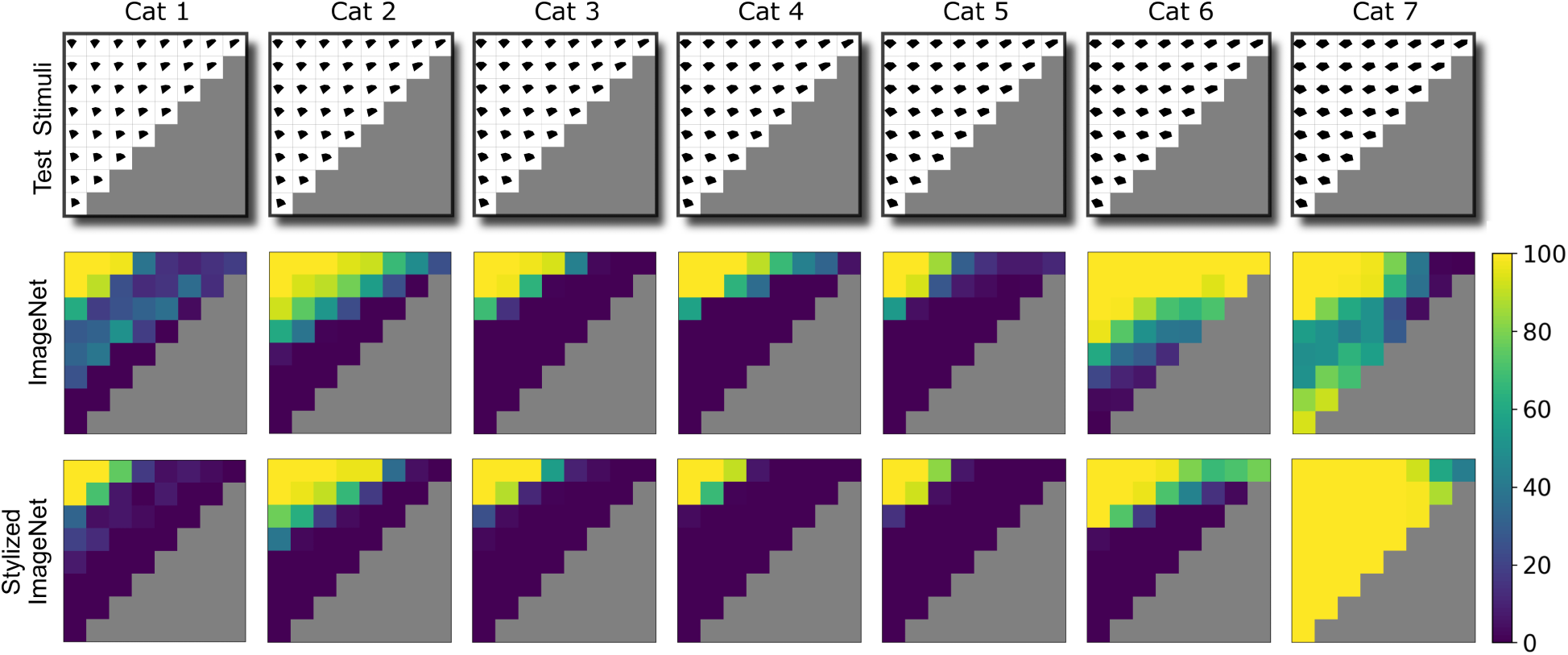
Performance of VGG-16 on deformations of single-part objects. *Note.* Test stimuli for each category is shown in the top row. Middle and bottom rows show accuracy on the landscape of relational and coordinate deformations for the network pre-trained on `ImageNet` (middle row) or `Stylized-ImageNet` (bottom row). In each case, the network was fine-tuned on the set of seven polygons shown in Figure 5(a). Each heatmap (in middle and bottom rows) corresponds to a category and shows the percent of shapes (with a relational and coordinate deformation given by the position on the landscape) accurately classified as the category from which the stimulus was derived. For most categories, accuracy is highest for small deformations (top-left corner) and decreases as a function of the coordinate distance from the basis shape (perpendicular to diagonal). The relational distance (left-to-right) has no added effect on this decrease in accuracy.

But classification accuracy only provides a indirect measure of internal representations. In order to get more insight into the network’s internal representations for relational and coordinate deformations, we examined the cosine distance between internal representations for the Basis image and two test images that were equidistant from it. An example of these images is highlighted (dashed red squares) in Figure 5(b). These cosine similarities for two `VGG-16` networks are plotted in Figure 7 (we again obtained qualitatively similar results for `AlexNet` – see Figure C6 in Appendix). At all internal layers, we observed that the average similarity between a Basis image and it’s relational (shear) deformation was equal or higher than the average similarity between the Basis image and it’s coordinate (rotation) deformation (compare solid (red) and dashed (blue) lines in Figure 7). In other words, relational deformation of an image was closer to the Basis image than it’s coordinate deformation and pre-training on the `Stylized-ImageNet` dataset to give the network a shape-bias did not change this pattern. This is the opposite of what one would expect if the network represented the stimuli in a relational manner.

**Figure 7.**
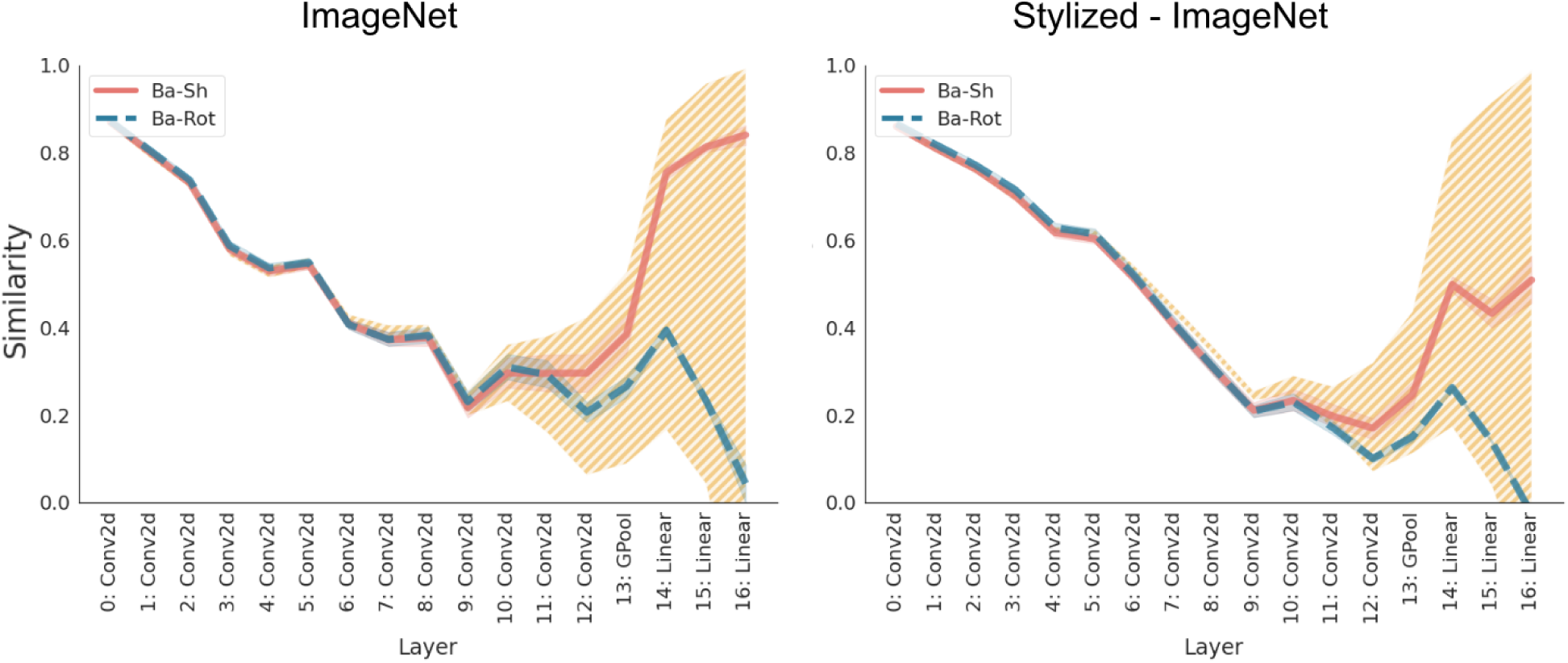
Cosine similarity in internal representations of VGG-16 in Experiment 3. *Note.* The solid (red) and dashed (blue) lines show the average cosine similarity between Basis images and relational (shear) and coordinate (rotation) deformations, respectively. The hatched (yellow) region shows the bounds on this similarity, with the upper bound determined by the average similarity between Basis images from the same category and lower bound determined by the average similarity between Basis images of different categories. If relational (shear) deformation has a larger affect on internal representations than a coordinate (rotation) deformation, one would expect the solid (red) line to be below the dashed (blue) line.

## Experiment 4

Results of Experiment 3 showed that a CNNs trained to classify objects do not show any enhanced sensitivity to deformations of relations between features of single-part objects. In other words, we did not observe any evidence suggesting that the CNNs infer properties of distal stimuli based on the proximal input image. In our next experiment, we examined how humans trained on the exact same stimuli responded to the two types of deformations.

### Participants

Participants (*N* = 37, *M_age_*= 33, 70% female, 30% male^2^) with normal or corrected-to-normal vision were recruited via Prolific for an online study and the experiment was conducted on the Pavlovia platform. They were reimbursed a fixed 2 GBP and participants who proceeded to the testing phase (*N* = 23) had a chance to earn a bonus of up to another 2 GBP depending on their performance during testing. The average payment was 8 GBP/hour. A written ethics approval for the study was obtained from the University of Bristol Ethics board.

### Stimuli

Four categories (out of seven) were chosen from the dataset in Experiment 3 to train participants. These were Cat 1, Cat 3, Cat 5 and Cat 7 from Figure 5(a). For the test data, we selected two deformations of each type that were matched according to the cosine distance from the basis (trained) image. For the relational deformation, these were the fifth (Deformation D1) and final (Deformation D2) shear in the top row of Figure 5(b). For the coordinate deformation, these were the fifth (D1) and final (D2) rotations in the left most column of Figure 5(b). This made up the 5 conditions in the experiment: Basis, D1 (Shear), D2 (Shear), D1 (Rotation) and D2 (Rotation). The original stimuli were 224×224 pixels but were re-scaled for each participant to 50% of the vertical resolution of the participant’s screen to account for the variability in screen size and resolution when running the study online.

### Procedure

Participants completed a supervised training phase in which they learned to categorize basis versions of the four categories. Each training block consisted of 40 stimuli for a total of 200 training trials (50 per category). Feedback on overall accuracy was given at the end of each block. Participants completed up to a maximum of 5 training blocks, or until they reached 85% categorization accuracy in a block. Participants who managed to reach 85% accuracy continued to the test block. The order of trials was randomised for each participant. Each trial started with a fixation cross (750 ms), then the stimulus was presented (500 ms) followed by four response buttons corresponding to the four categories (until response). After participants responded, feedback was given - CORRECT (1 s) if the response was correct, and INCORRECT with additional information about what the correct response should have been (1.5 s) if the response was incorrect. The training phase was followed by a test phase consisting of five test blocks. Each block consisted of 20 trials for a total of 100 test trials (25 per condition). Like the training phase, the order of test trials was randomised for each participant. The procedure for each test trial was the same as in the training phase except that participants were not given any feedback during testing.

### Analysis

Four planned comparisons (t-tests) were conducted in order to test whether accuracy rates in each of the shear and rotation conditions differed from accuracy in the basis condition.

### Results and Discussion

The average CNN and human accuracy of classification on each of these deformations is shown in Figure 8. We can see that irrespective of training, `VGG-16` was more sensitive to rotation than to shear (see Figure C7 for `AlexNet`). While performance decreases for both deformations, it decreases more rapidly for rotations. Human participants showed the opposite pattern (Figure 8, right-hand panel). There was no significant difference in performance between the basis image and the two rotation deformations (both *t*(22) *<* 3.48, *p > .*28), while performance decreased significantly for each of the shear deformations (both *t*(22) *>* 14.10, *p < .*001, *d_z_ > .*83). The largest shear resulted in largest decrease in performance (*M_difference_* = 25.87%). Thus, the behaviour of participants was in line with the prediction of structural description theories, where shape is encoded based on relations between features, and in the opposite direction to the performance of the CNNs trained to classify objects.

**Figure 8.**
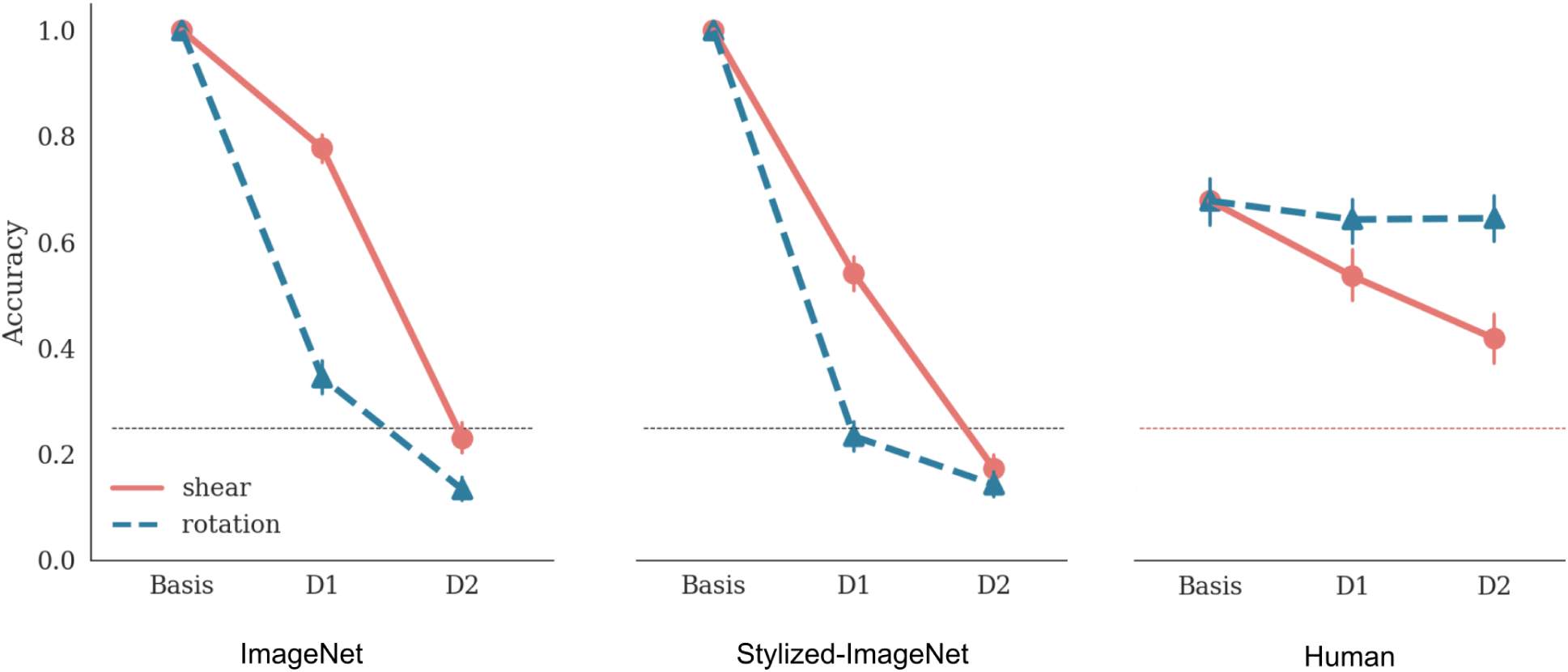
Comparison of humans and VGG-16 on how classification accuracy changes with deformations. *Note.* Left and middle panels show classification accuracy for `VGG-16` trained on `ImageNet` and `Stylized-ImageNet`, respectively. Right panel shows performance of human participants on the same stimuli. Each panel shows performance under three conditions: basis image, deformation D1 and deformation D2. For the shear deformation (solid, red line), D1 and D2 consists of images in the top row in the fourth and eighth column in Figure 5. For the rotation deformation (dashed, blue line), D1 and D2 consist of images in the first column and fourth and eighth rows. Error bars show 95% confidence interval and dotted black line shows chance performance.

## Experiment 5

One response to the difference between CNNs and humans in Experiments 3 and 4 is that it arises due to the difference in experience of the two systems. Humans experience objects in a variety of rotations and consequently represent a novel object in a rotation invariant manner. CNNs, on the other hand, have not been explicitly trained on objects in different orientations (although `ImageNet` includes some objects in various poses). It could therefore be argued that CNNs do not learn relational representations in Experiment 3 because the training set did not provide an incentive for learning such a representation. Indeed, the optimisation view argues that a bias must be present in the training environment for the visual system to internalise it.

To give the network a better chance of learning to classify based on internal relations, we conducted two further simulations. In the first simulation, we trained the networks on some rotations for all Basis shapes and tested them on unseen rotations. This simulation emulates generalising the concept of rotation for each object after observing some of the rotations for that object. In the second simulation, the networks were shown *all* rotations of some Basis shapes and tested on unseen rotations of the left-out Basis shapes. This simulation emulates generalising the concept of rotation from one object to another.

### Methods

All methods in Experiment 5 remained the same as Experiment 3, except for the images in the training sets. In the first simulation, the training set now consisted of Basis (polygon) shapes presented at random translations and scales (just like Experiment 3) but additionally, also at rotations in the range [*−*45*^◦^,* 0] for all polygons. We then tested the networks on rotations in the range [0, +45*^◦^*]. In the second simulation, we selected six (out of seven) categories and trained the network on random translations, scales and *all* rotations ([0, 360°)) for these categories. For the seventh category (Cat 3), images were still randomly translated and scaled, but always presented in the upright orientation. We then tested how the network generalised to the two types of deformations for this critical category. We obtained qualitatively similar results for networks pre-trained on `ImageNet` and `Stylized-ImageNet`. Since the network trained on `Stylized-ImageNet` has the best chance of capturing human data, here we present the results of this network for both simulations.

### Results and Discussion

The network performance for the first simulation is shown in Figure 9. We observed that, despite being trained on this augmented dataset, results remained qualitatively similar. For most categories performance degraded equally or more with a change in rotation than with an equivalent change in shear. That is, the network was *better* at generalising to large relational deformations (shears) than large relation-preserving deformations (rotation). The pattern was different for Category 6, where the network showed good performance on large rotations. But examining the confusion matrix again revealed that the high accuracy at large rotations for this category was misleading as it was accompanied with large Type I errors: large rotations for shapes of any category were mis-classified as belonging to the Category 6. Overall, we did not find any evidence for the network learning shapes based on their internal relations.

**Figure 9.**
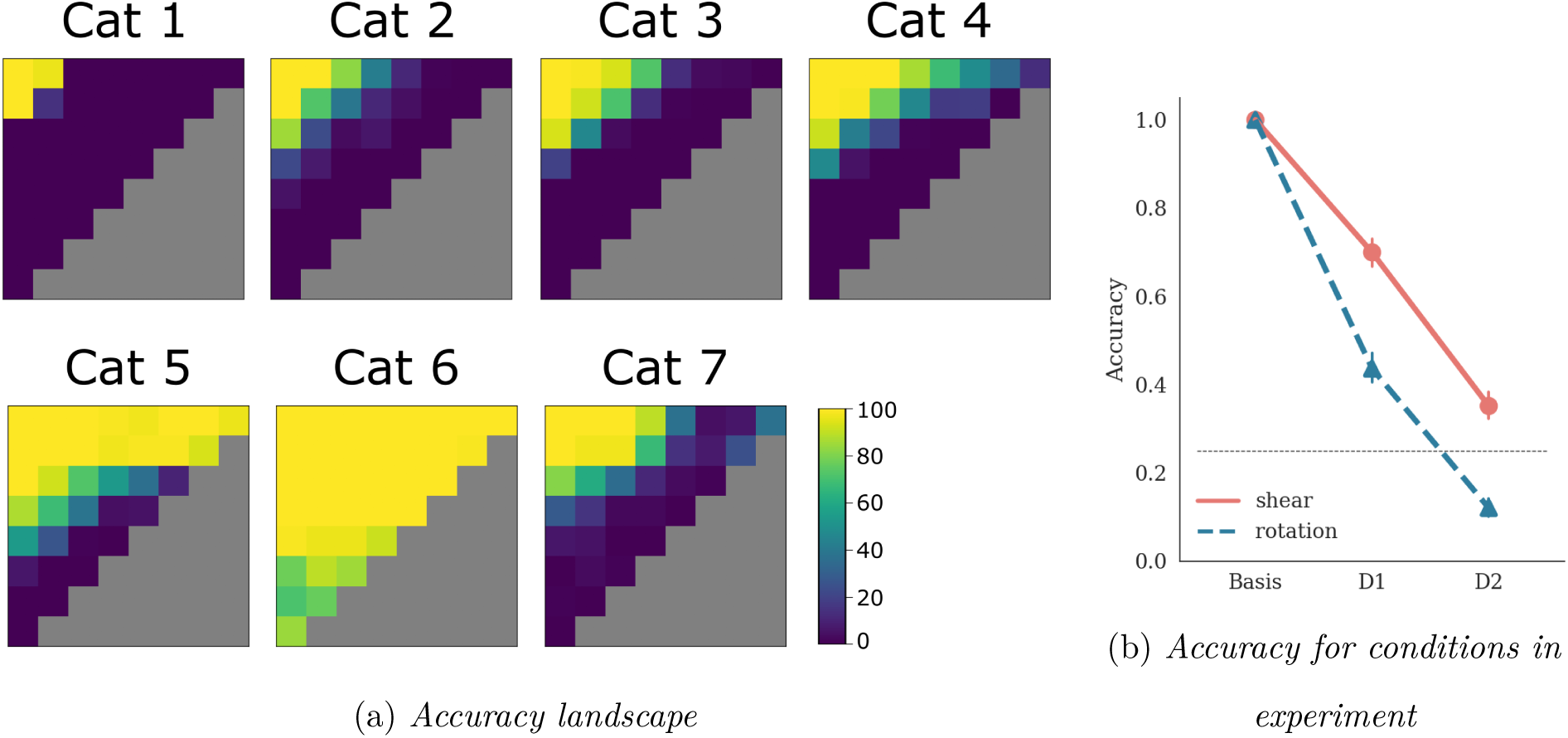
Performance of VGG-16 trained on some rotations of all categories. *Note.* (a) Accuracy of network plotted as percent of correct classifications for each rotation and shear deformation of each category, and (b) accuracy for shear (solid, red) and rotation (dashed, blue) as a function of deformations used in Experiment 4 with human participants (compare with Figure 8, right-hand panel).

The results of the second simulation are shown in Figure 10. Figure 10a shows the heat-map of accuracy on the test grid for the left-out category. This heat map showed that the network continued showing the pattern observed above – it’s performance decreases across (perpendicular to) the diagonals, but increases as one moves from left-to-right along these diagonals. Figure 10b shows the performance on the same conditions as the human experiment (see Figure 8). Again, we see that the performance drops less rapidly across the two shear deformations (dashed line) than the two rotation deformations (solid line). This figure makes it clear that training other orientations on all rotations does not help the network generalise better to novel orientations for the left-out category. In fact, the performance drops more quickly than when none of the categories were rotated in the training set (compare with Figure 8). This is because the network starts classifying novel orientations of the left-out shape as the shapes that it had seen being rotated in the training set.

**Figure 10.**
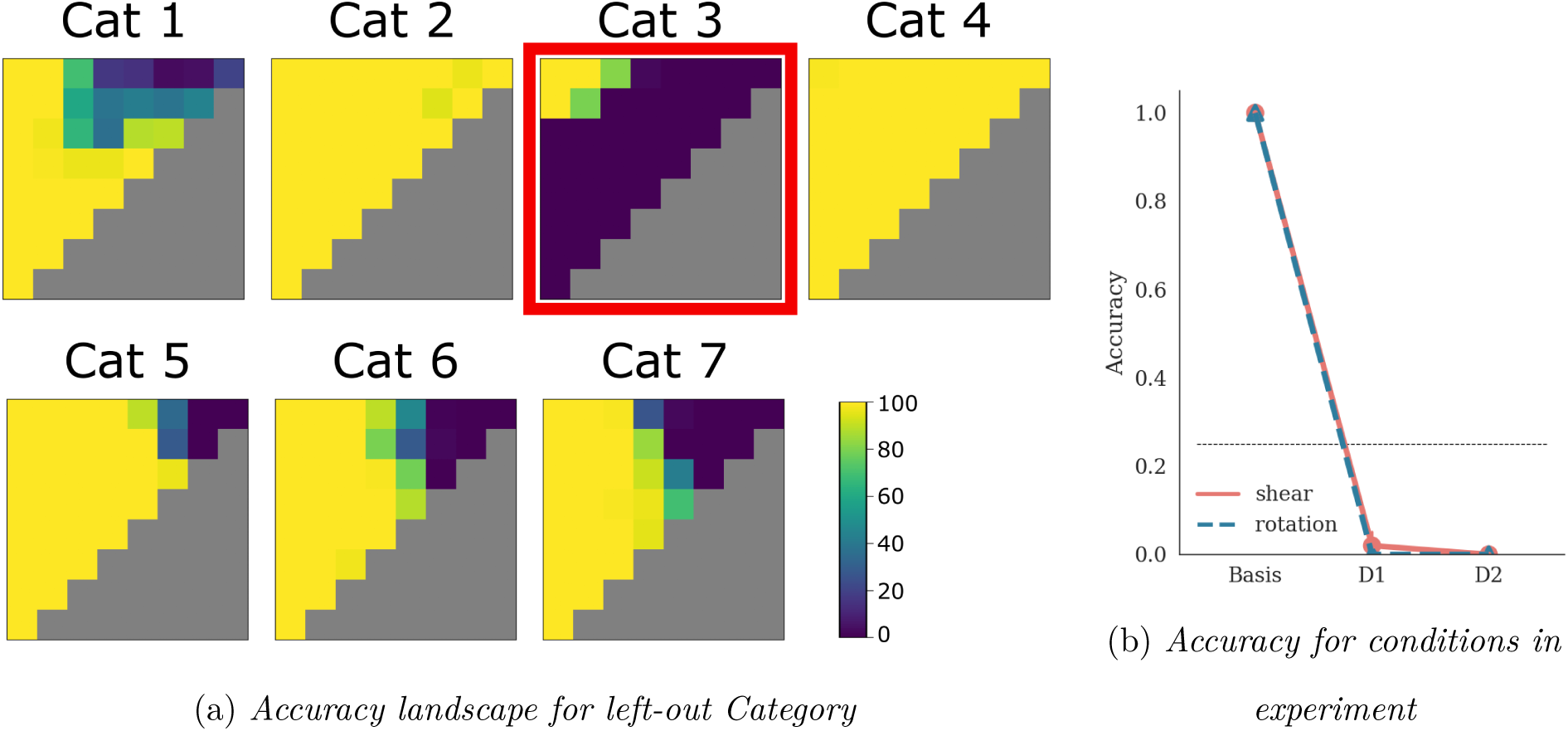
Performance of VGG-16 trained on all rotations of some categories. *Note.* (a) Accuracy of network plotted as percent of correct classifications for rotation and shear deformations for all categories. Note the high performance for all rotations of most categories is expected as the network is trained on these rotations. The critical (out-of-training-distribution) test is the network’s performance on the left-out category – Category 3 (highlighted using red rectangle) (b) Accuracy of network for the set of deformations D1 and D2 for Category 3, tested in Experiment 4 with human participants (compare with Figure 8, right-hand panel).

It may be tempting to think that the differences between humans and CNNs can be reconciled by training CNNs that learn rotation-invariant shapes. However, consider how a CNN achieves rotation-invariance. Figure 11, taken from Goodfellow, Bengio, and Courville (2016, chap. 9), illustrates how a network consisting of convolution and pooling layers may learn to recognise digits in different orientations. As a result of training on digits (here, the digit 5) oriented in three different directions, the convolution layer develops three different filters, one for each orientation. A downstream pooling unit then amalgamates this knowledge and fires when any one of the convolution filters is activated. Therefore, this pooling unit can be considered as representing the rotation-invariant digit 5. During testing, when the network is presented the digit 5 in any orientation, the corresponding convolution filter gets activated, resulting in a large response in the pooling unit and the network successfully recognises the digit 5, irrespective of it’s orientation.

**Figure 11.**
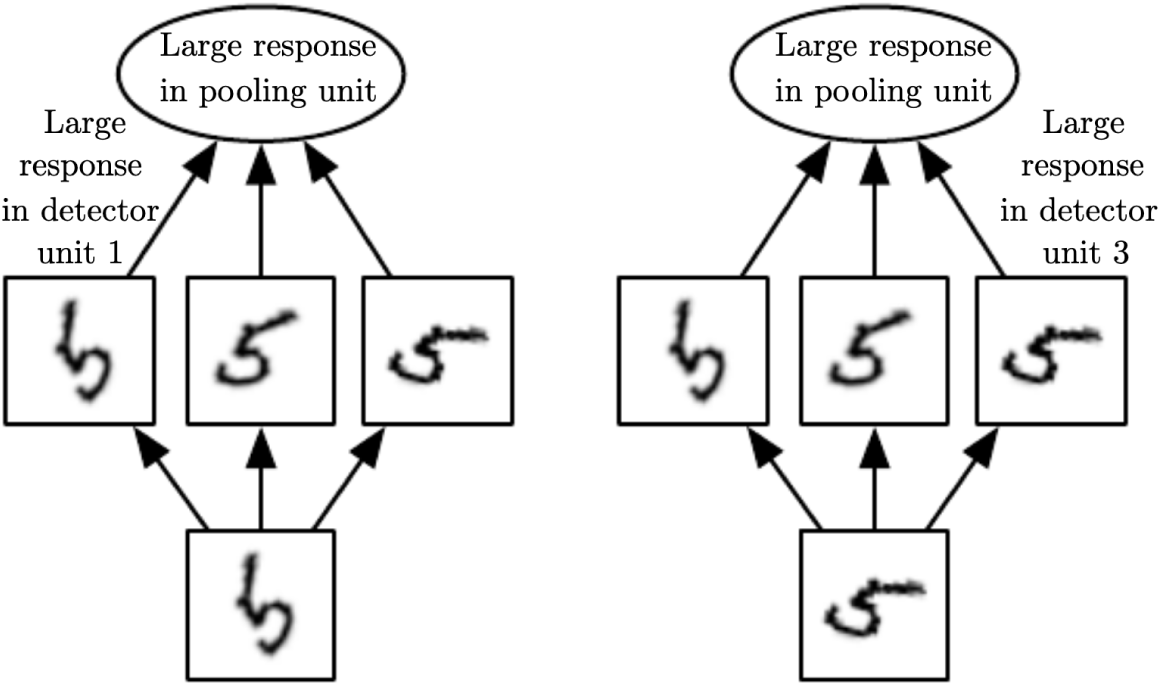
A proposal for achieving rotation invariance in CNNs (from Goodfellow et al. (2016, chap. 9)) *Note.* Each panel shows the response of the network to a different rotation of the digit ‘5’. The network detects different rotations by having a large set of filters, one matching each rotation. The output unit pools across all rotation filters, essentially performing a disjunction over all filter activations.

In contrast, a relational account of shape representation does not rely on developing filters for each orientation of a shape. Indeed, it is not even necessary to observe a shape in all orientations to get, at least some degree of, rotation invariance. All that is needed is to be able to recognise the internal parts of an object and check whether they are in the same relation as the learned shape. Accordingly, many psychological studies have shown that invariance, such as rotation invariance, precedes recognition (Biederman & Cooper, 1991, 1992; Biederman & Gerhardstein, 1995; Hummel, 2013).

## General Discussion

In a series of experiments we have shown that humans represent shape in qualitatively different ways to CNNs that learn to classify large datasets of objects using supervised learning. In Experiment 1 we found that CNNs trained to classify objects were entirely insensitive to deformations in categorical relations between object parts. Furthermore, we could not train CNNs to be sensitive to relational changes in general even when we made relational changes diagnostic of category classification (Experiment 2). In Experiment 3 and 4, where we precisely matched the extent of relational and coordinate deformations, we found that humans were highly sensitive to relational deformations of single-part objects, whereas CNNs were only sensitive to coordinate distance, and once again, CNNs could not learn to be sensitive to relational manipulations (Experiment 5).

These findings challenge the hypothesis that humans perceive objects based on similar principles as CNNs trained to classify large sets of objects and that apparent differences arise due to “differences in the data that they see” (Hermann et al., 2020). These results show that even CNNs that have been trained to classify objects on the basis of shape (trained on the `Stylized-ImageNet`) learn the wrong sort of shape representation. These findings add to other studies that also highlight the different types of shape representation used by CNNs and the human visual system. For example, Puebla and Bowers (2021) have found that CNNs fail to support a simple relational judgement with shapes, namely, whether two shapes are the same or different. Again, this highlights how CNNs trained to process shape ignore relational information. In addition, Baker et al. (2018) have shown that CNNs that classify objects based on shape focus on local features and ignore how local features relate to one another in order to encode the global structure of objects.

These failures may reflect a range of processes present in humans but absent in CNNs trained to recognise objects through supervised learning, such as figure-ground segregation, completing objects behind occluders, encoding border ownership, and inferring 3D properties about the object (Pizlo et al., 2010). Consistent with this hypothesis, Jacob, Pramod, Katti, and Arun (2021) and Bowers et al. (2022) have recently highlighted a number of these failures in CNNs, including a failure to represent 3D structure, occlusion, and parts of objects. More broadly, these results challenge the the claim that CNNs trained to recognise objects through supervised learning are good models of the ventral visual stream of human vision (see, for example, (Cadieu et al., 2014; Mehrer, Spoerer, Jones, Kriegeskorte, & Kietzmann, 2021; Yamins et al., 2014)).

One interesting study that provides some evidence to suggest that standard CNNs have similar shape representations to humans was reported by Kubilius, Bracci, and Op de Beeck (2016). In one of their experiments (Experiment 3), they compared the similarity of representations in various CNNs in response to a change in metric and non-accidental features of single-part objects. For instance, they compared a base object that looked like a slightly curved brick to two objects: one object that was obtained by deforming the base object into a straight brick (a non-accidental change) and a second object that was obtained by deforming the base object into a greatly curved brick (a metric change). Kubilius et al. reported that, like humans, CNNs were more sensitive to non-accidental changes. However, it is unclear whether CNNs were more sensitive to one of their manipulations because of the non-accidental change or because of other confounds accompanying these manipulations. For example, when Kubilius et al. modified some of the base shapes to non-accidental deformations, it was accompanied by a change in local features (such as properties of vertices). Recent research (Baker et al., 2018; Geirhos et al., 2018) has shown that, unlike humans, CNNs are in fact highly sensitive to change in local and textural features and it is unclear whether it is these types changes that are driving the effects observed by Kubilius et al. (2016). More work is required to reconcile their findings with our own.

More generally, our findings raise the question as to whether optimizing CNNs on classification tasks is even the right approach to developing models of human object recognition. It is striking how well our findings are well predicted by a classic structural description theory of object recognition that builds a distal representation of objects using heuristics (e.g., Biederman, 1987). As detailed above, on this theory, the visual system encodes specific features of the proximal stimulus that are best suited for making inferences about the distal object. This includes explicitly coding the relations between parts in order to support visual reasoning about objects (e.g., appreciating the similarity and differences of buckets and mugs as discussed above), and encoding parts in terms of non-accidental features that often include relations between features, such as symmetry, in order to infer their 3D distal shape from variable proximal 2D images. Just as predicted, humans are selectively sensitive to these deformations (changes in the relations between parts in Figure 1 and changes in symmetry in Figure 5), whereas CNNs treated these deformations no differently than others.

Of course, it is possible that training CNNs on a range of different tasks (especially tasks where the objective is to approximate the distal representation) or on tasks with different objectives rather than classification (e.g. unsupervised learning of image sequences (Parker & Serre, 2015), or generative modelling (Kingma & Welling, 2013) or on a “self-supervised” task (Grill et al., 2020)) may lead to shape representations that are more similar to those formed in human visual cortex. However, here we wanted to focus on CNNs trained on recognising objects through supervised learning because of two reasons. Firstly, it has been argued that CNNs trained under these settings learn to classify objects based on human-like shape representations (Geirhos et al., 2018; Hermann et al., 2020; Kubilius et al., 2016). Secondly, these models have had the largest success in predicting neural representations in human and primate visual system (Cadieu et al., 2014; Schrimpf et al., 2020; Yamins & DiCarlo, 2016) and it has been argued that there is a “strong correlation between a model’s categorization performance and it’s ability to predict individual level IT neural unit response data” (Yamins et al., 2014). Our findings challenge the view that optimizing performance in a classification task can explain shape representations used during human shape perception. Instead, these findings are well predicted by the classic structural description theory of object recognition that builds a distal representation of objects using heuristics (e.g., Biederman, 1987).

It is also possible that a different Deep Learning architecture may be more successful than CNNs at encoding objects based on relations between their parts. Indeed, previous research indicates that relational reasoning may require a more powerful architecture that can explicitly and separately represent (i) parts and relations, and (ii) their bindings (e.g., to distinguish whether the brick is above the cone or vice-versa; Doumas, Puebla, Martin, & Hummel, in press; Hummel, 2011; Hummel & Biederman, 1992; Hummel & Holyoak, 1997, 2003). Other Deep Learning architectures such as Capsule Networks (Sabour, Frosst, & Hinton, 2017), Transformers (Vaswani et al., 2017), LSTMs (Hochreiter & Schmidhuber, 1997) or Neural Turing machines (Graves, Wayne, & Danihelka, 2014) may also provide the representational power necessary to represent structural descriptions. What is clear from our study is that learning to classify objects is not, in and of itself, sufficient for the emergence of human shape representations.

## Code and Data

All code for generating the datasets, simulating the model as well as participant data from Experiment 4 can be downloaded from: https://github.com/gammagit/distal

## Appendix A Classification performance

**Figure A1.**
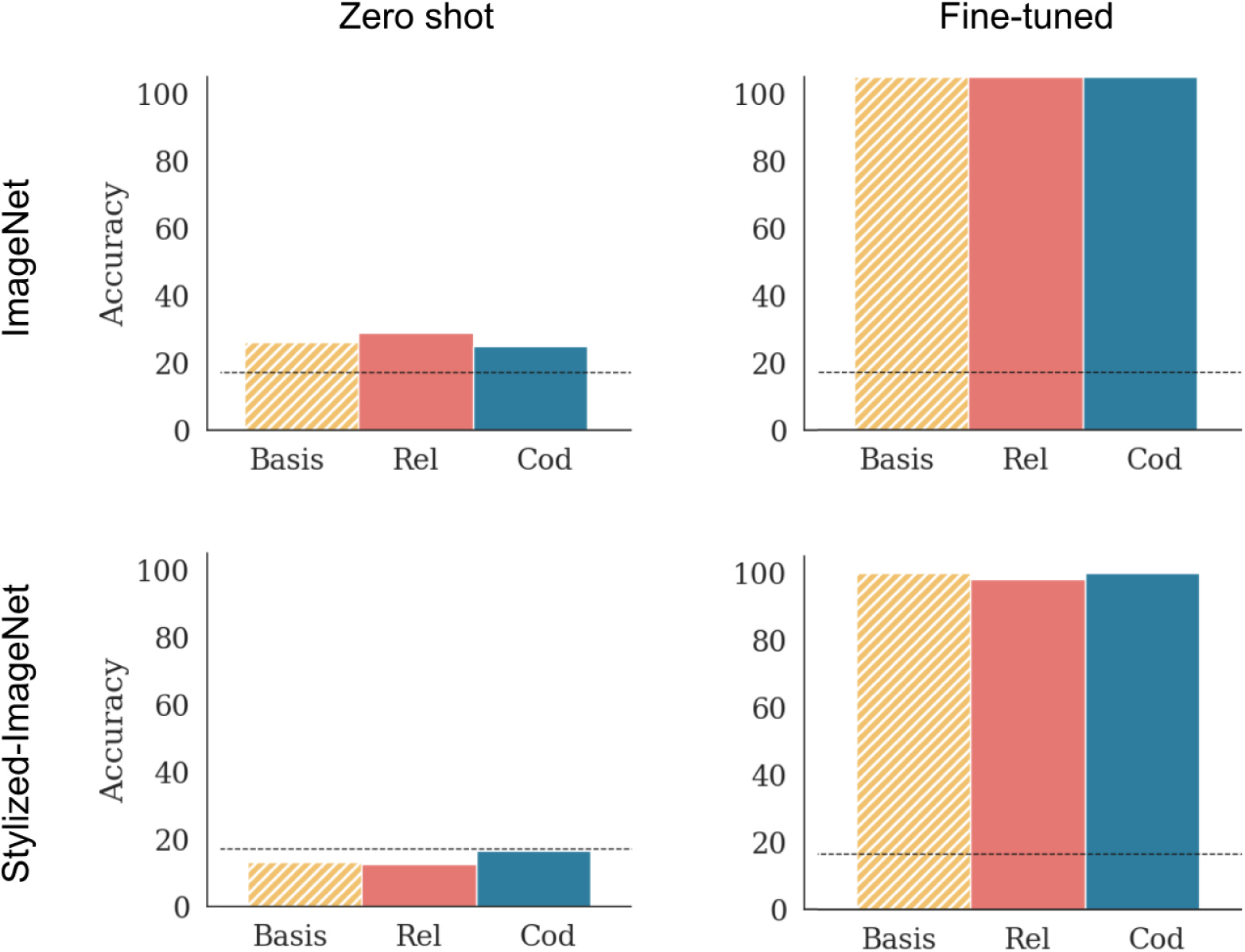
Classification accuracy for VGG-16 in Experiment 1. *Note.* Panels in the first column show classification accuracy on the pre-trained network without any further training, while panels in the second column shown test performance for a model that was fine-tuned on the set of Basis shapes. Dashed black line shows chance performance. In keeping with the cosine distance in internal representations, we observed that models in the *Zero-shot* condition failed to classify the Basis shapes or their deformations (accuracy was statistically at chance across models) and models in the *Fine-tuned* condition learned to perfectly classify Basis images, but failed to distinguish them from relational or coordinate deformations.

## Appendix B Examining errors in Experiment 3

**Figure B1.**
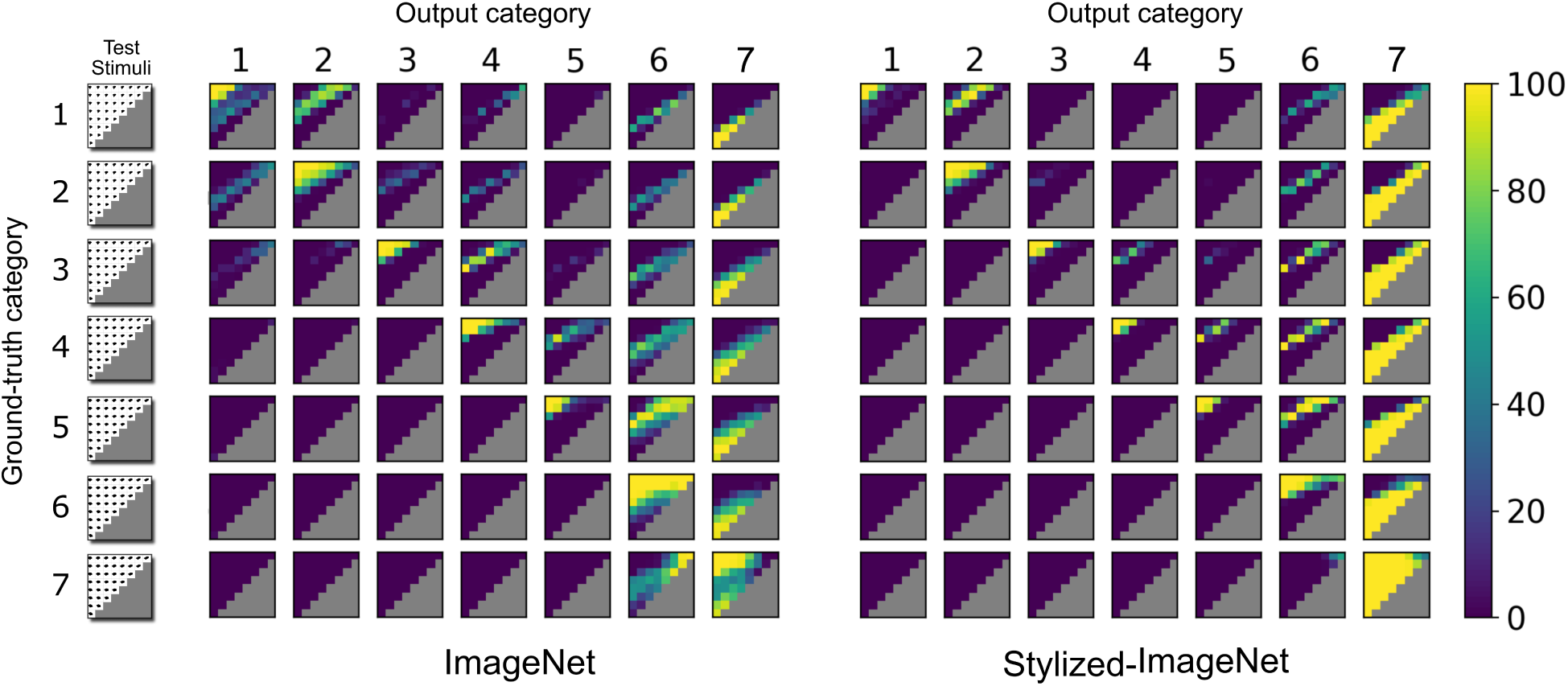
Confusion matrices for VGG-16 in Experiment 3. *Note.* For any heat map, the category label along each row shows the ground truth – i.e., all test shapes used to obtain the heat map were obtained by distorting the basis shape from that category. The category label along the column shows output category label assigned by the network. Therefore, in each row, the diagonal heat map shows the correct classifications, while the off-diagnoal heat maps show how each deformation was misclassified.

In Experiment 3, we observed that performance decreased as a function of coordinate distance for most categories. However, in most simulations, we also observed that there was one category where performance was really high for most deformations, including large rotations. For example, in Figure 6, most categories show a large decrease in performance with increase in rotation of test images, except for Category 7 (both middle and bottom rows). To understand why this was the case, it is useful to look at the errors made by the network. Figure B1 shows confusion matrices for two models (`VGG-16` pre-trained on `ImageNet` and `Stylized-ImageNet` respectively). Each heat map shows the number of times an output category was chosen for all deformations of a given input category. This confusion matrix shows that the both networks were prone to mis-classify large rotations from any category as belonging to Category 7 (note large number of classifications in final column of each matrix for large rotations). These false positives (Type I errors) create a bias in the accuracy results for Category 7 in Figure 6 – that is, the high accuracy for large rotations for Category 7 category are, in fact, misleading as the networks classify large rotations for any category as Category 7. These confusion matrices also show that the networks showed a “rightward” bias – there are more Type I errors in the upper triangle of each matrix than the lower triangle. In other words, the network was more likely to mis-classify images from each category as the category above rather than the category below.

## Appendix C Results for AlexNet

**Figure C1.**
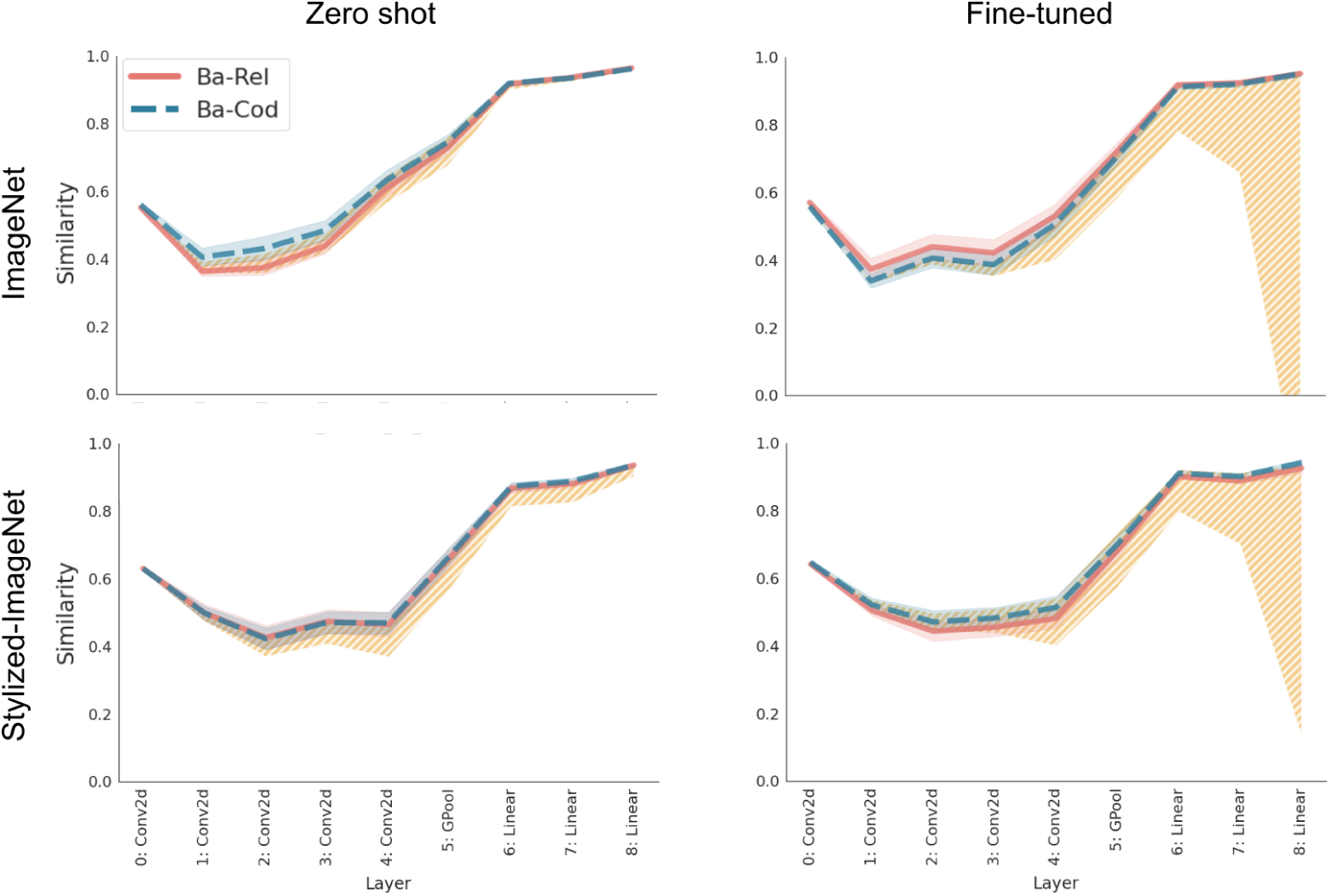
Cosine similarity for internal representations for Alexnet in Experiment 1. *Note.* Like the results for `VGG-16` (compare with Figure 2 in the main text), the similarity between Basis images and both types of deformations is at the upper bound throughout the network, showing that the network does not distinguish the trained (Basis) image from it’s Rel and Cood deformations.

**Figure C2.**
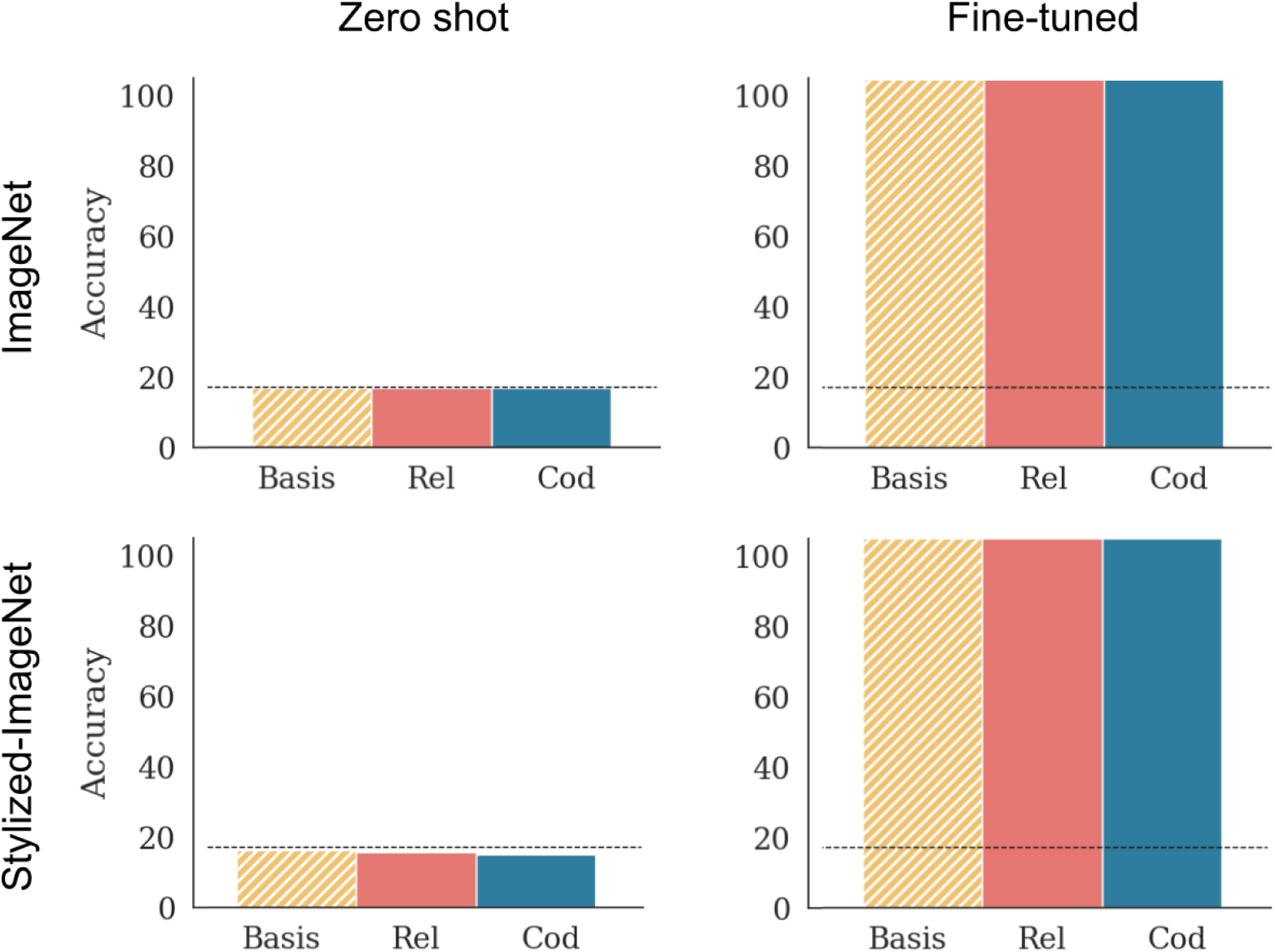
Performance of AlexNet in the test set for Experiment 1. *Note.* Each panel shows accuracy on the Basis shapes as well as the two types of deformations: relational (Rel) which changes a categorical relation and coordinate (Cood), which preserves all categorical relations. Compare with performance of `VGG-16` in Figure A1.

**Figure C3.**
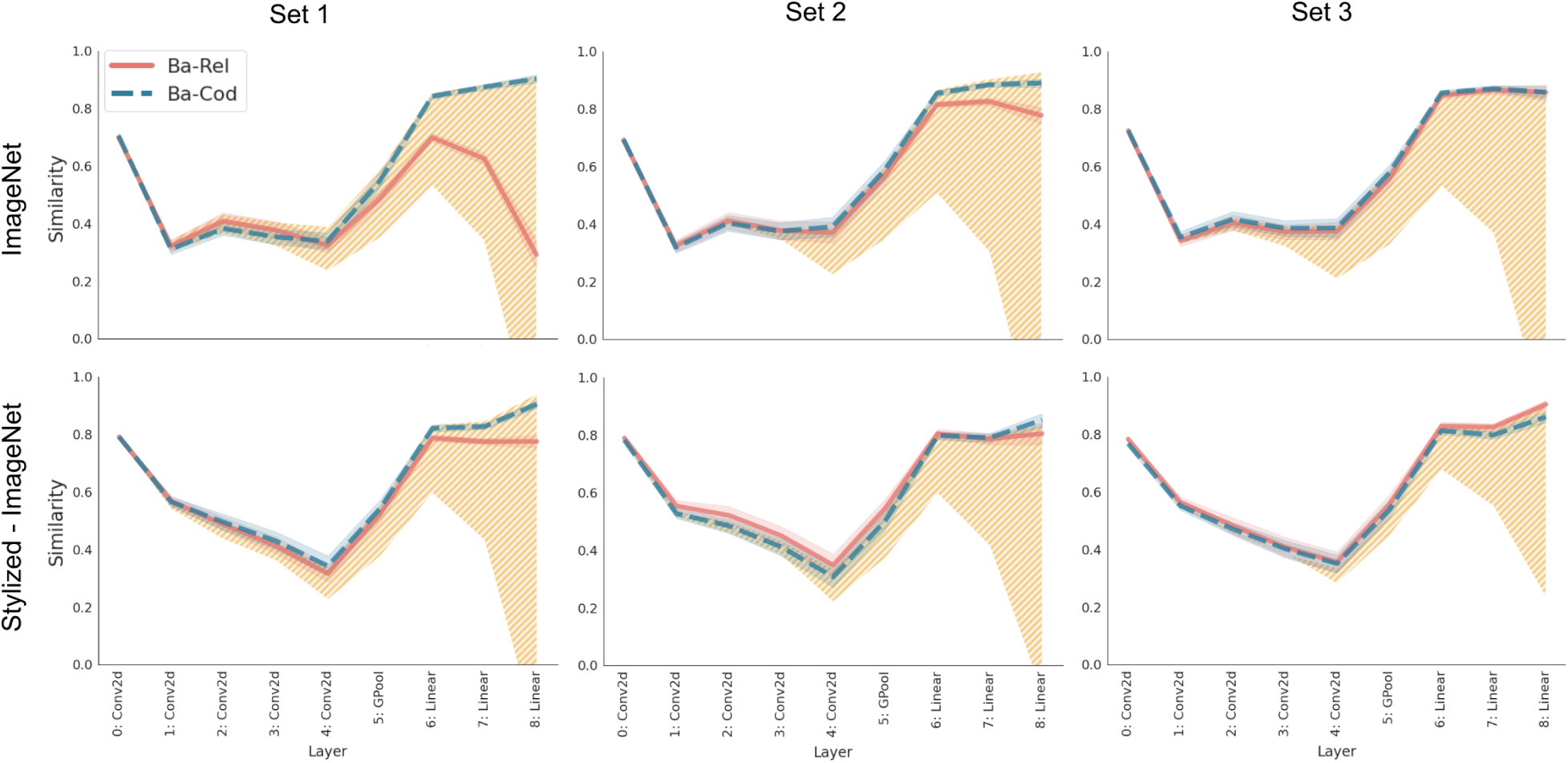
Cosine similarity for AlexNet trained on diagnostic relations in Experiment 2. *Note.* Like the results for `VGG-16` (compare with Figure 4), we see that networks learns to distinguish the Rel deformation from the Basis image for Set 1 (left column), when it has seen the specific deformation in the training set. But this sensitivity to Rel deformation diminishes in Set 2 (middle column), when only one pair of trained shapes have a similar deformation and completely lost for Set 3 (right column) when the network has been trained on the Rel deformations, but the specific deformation tested is novel.

**Figure C4.**
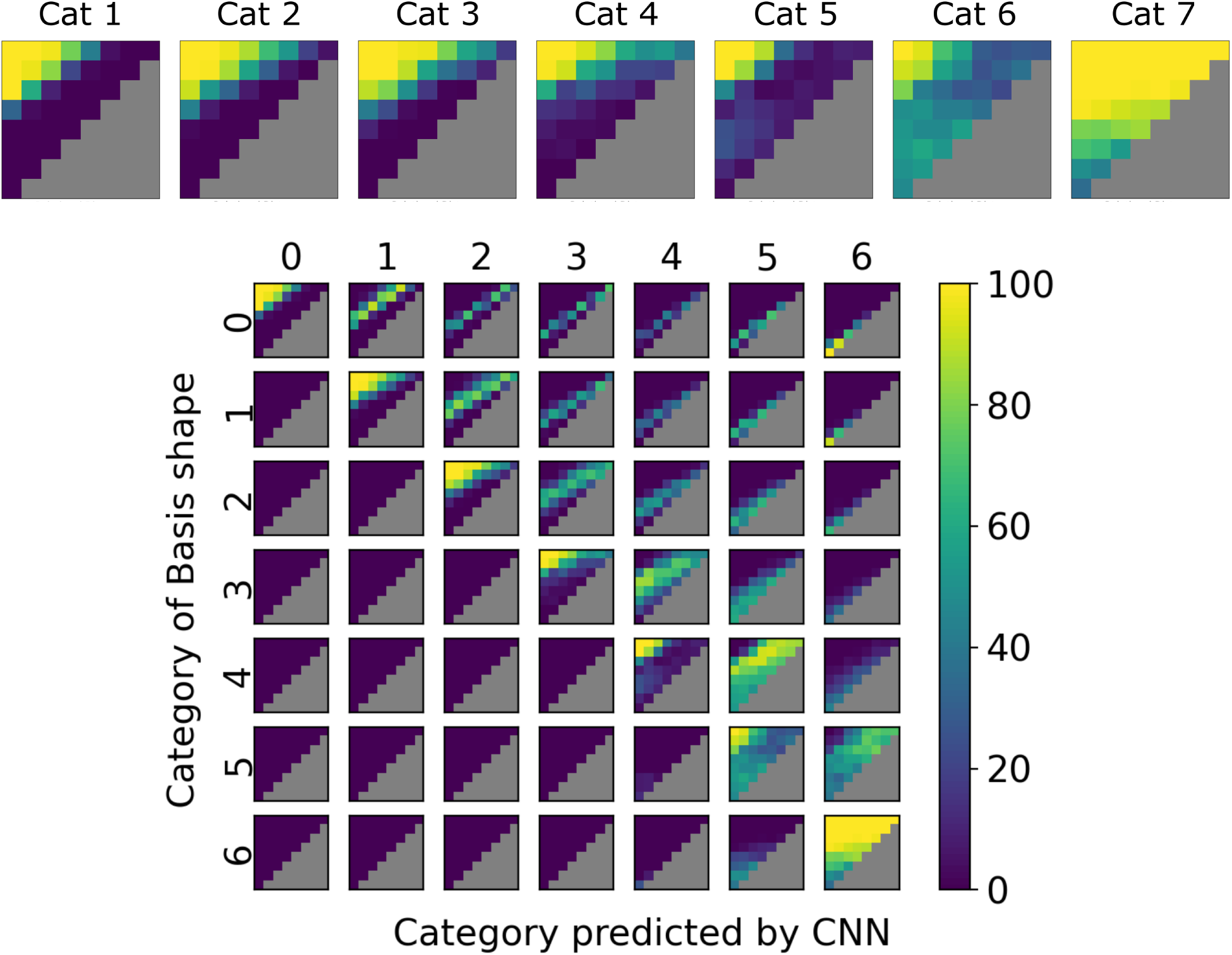
Classification performance for AlexNet trained on ImageNet in Experiment 3. *Note.* Each heatmap shows accuracy for on test items for a particular category for `AlexNet` pre-trained on `ImageNet` and fine-tuned on the dataset in Figure 5. Each cell in the heatmap corresponds to a deformation that is a combination of relational (shear) and coordinate (rotation) transformations of the trained Basis shapes (see Figure 5(a)). The grid at the bottom shows the “confusion matrix” – each heatmap in the grid shows the proportion of responses predicted as the category along the column for a deformation with basis shape taken from the category along the row. Like the results for `VGG-16` (compare with Figure 6), we see that accuracy decreases as a function of coordinate distance from the basis shape, rather than the relational distance.

**Figure C5.**
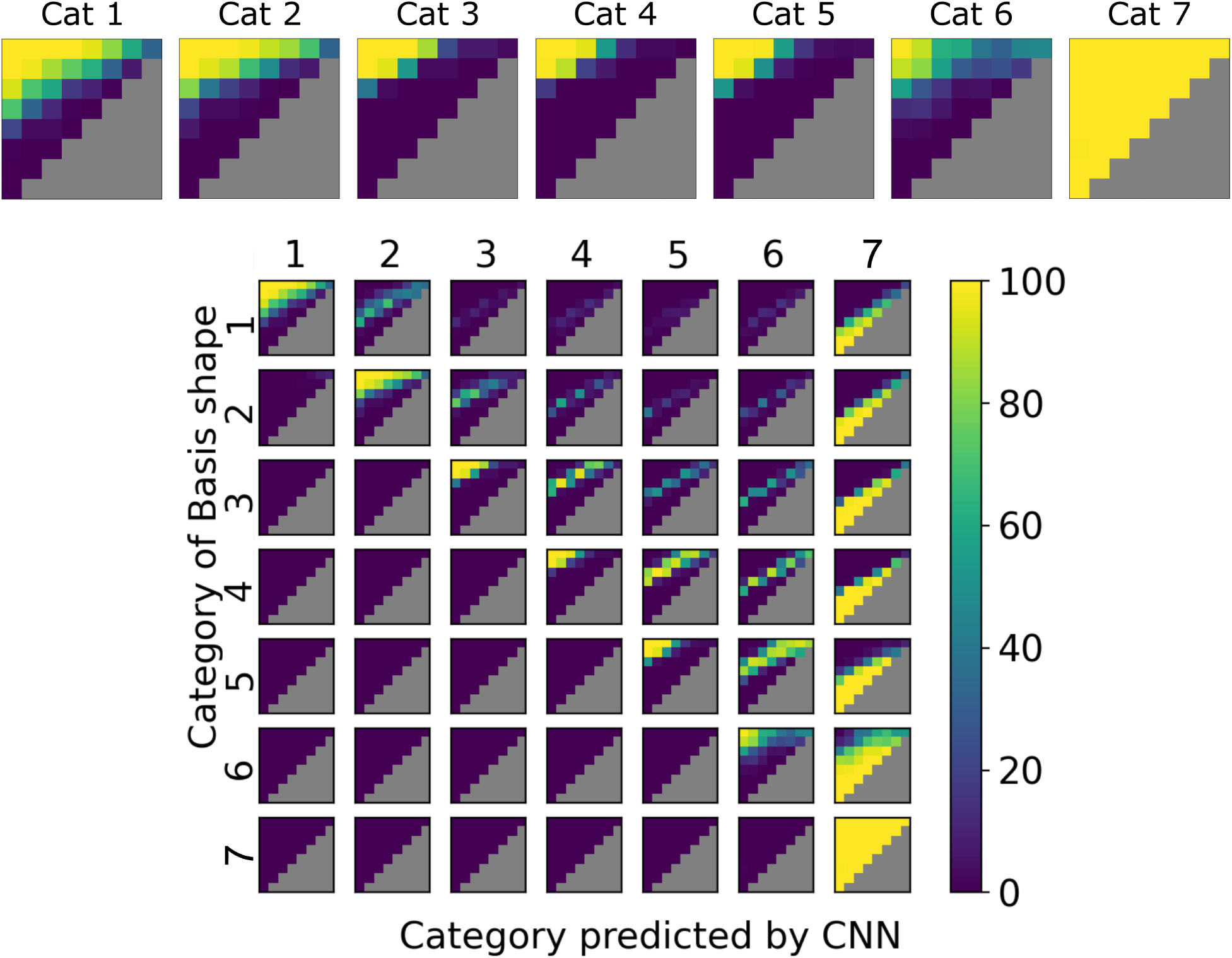
Classification performance for AlexNet trained on Stylized-ImageNet in Experiment 3. *Note.* Each heatmap in the top row shows accuracy for on test items for a particular category for `AlexNet` pre-trained on `Stylized-ImageNet` and fine-tuned on the dataset in Figure 5. The bottom panel shows the confusion matrix. See Figure C4 for explanation.

**Figure C6.**
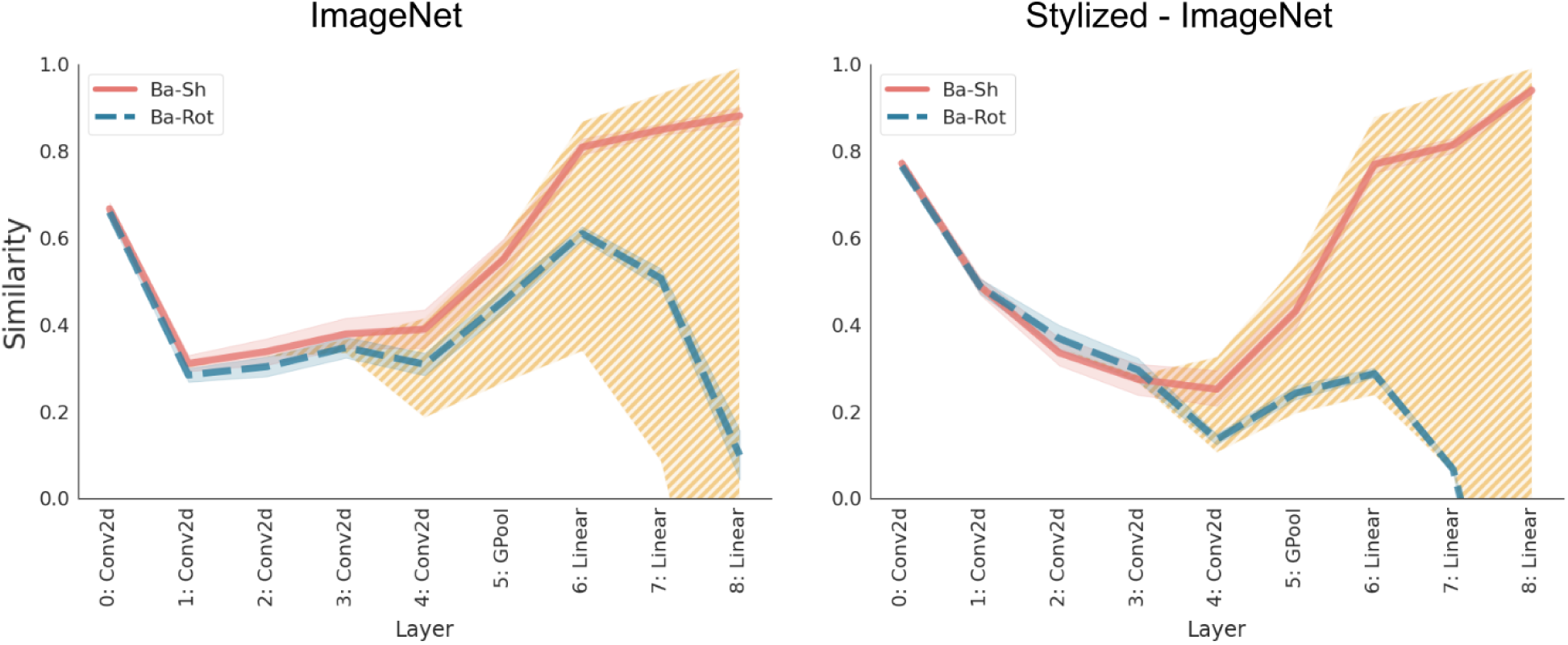
Cosine similarity for AlexNet in Experiment 3. *Note.* Cosine similarity between internal representations for the Basis shapes and two deformations of the basis shape (dashed red squares in Figure 5(b)) from the polygons dataset at each convolution and fully connected layer of `AlexNet`. Solid (red) line shows the average similarity between representations for a basis shape and its relational (shear) deformation, while dashed (blue) line shows the average similarity between a basis shape and it’s coordinate (rotation) transformation. The hatched area shows the bounds on similarity, with the upper bound determined by the average similarity between two basis shapes from the same category and lower bounds determined by the average similarity between two basis shapes of different categories. Like the results for `VGG-16` (compare with Figure 7), we observed that the network treated the relational (shear) deformation as being more similar to the basis shape than the coordinate (rotation) deformation.

**Figure C7.**
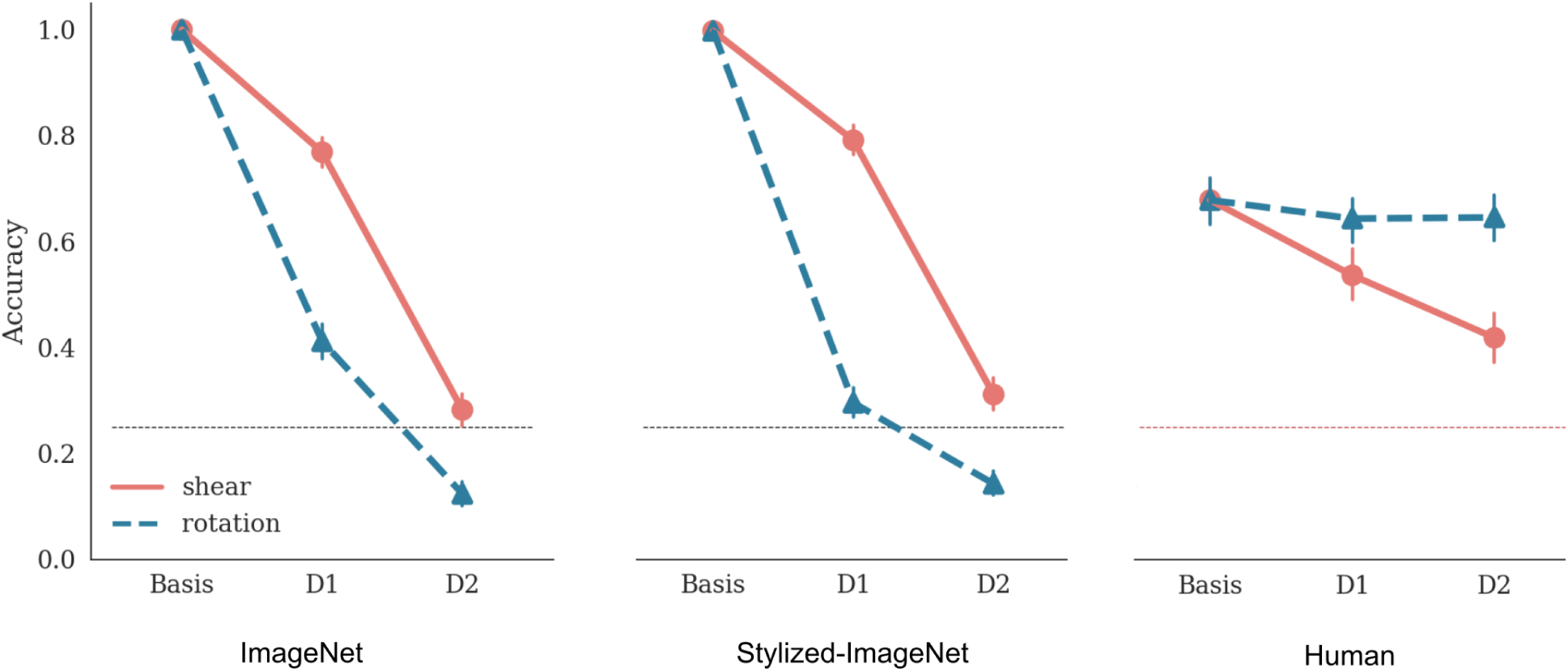
Comparison of classification accuracy for AlexNet (Experiment 3) and human participants (Experiment 4). *Note.* Each panel shows performance under three conditions: basis image, deformation D1 and deformation D2. For the shear deformation (solid, red line), D1 and D2 consists of images in the top row in the fourth and eighth column in Figure 5. For the rotation deformation (dashed, blue line), D1 and D2 consist of images in the first column and fourth and eighth rows. Error bars show 95% confidence interval and dashed red line shows chance performance. Note that the results in right-hand panel are reproduced here for convenience but are the results of the same experiment reported in Figure 8, right-hand panel.

**Figure C8.**
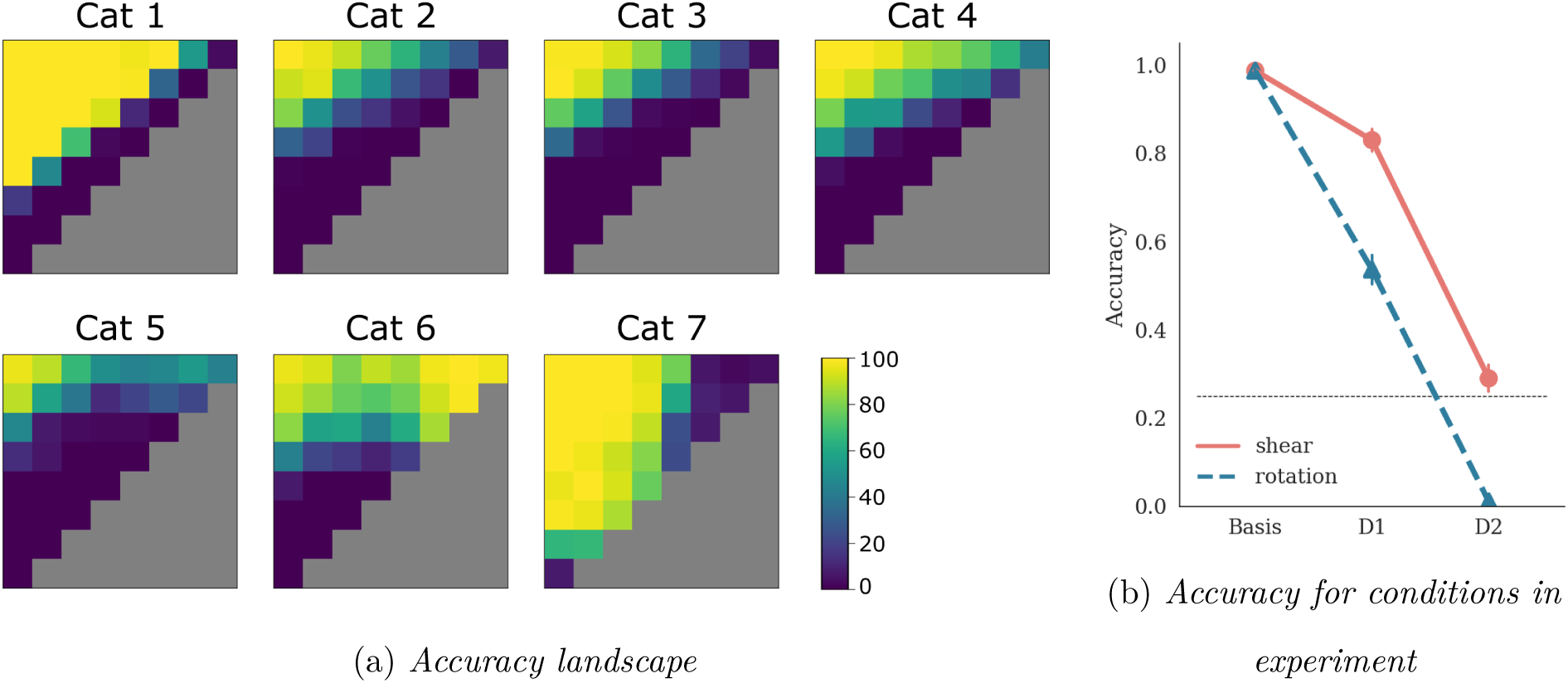
Performance of AlexNet in Experiment 5, Simulation 1. *Note.* Performance of `AlexNet` fine-tuned on an augmented dataset where the Basis shapes are not only translated and scaled but also randomly rotated in the range [*−*45°, 0°]. The network is then tested on on shear and rotation deformations in the range [0°, +45°]. Like the results for `VGG-16` (compare with Figure 9), we observed that even when the network was trained on some rotations, it’s performance on untrained rotations (a coordinate transformation) was still worse than shears (a relational transformation). (b) shows accuracy for deformation level D1 and D2 used for testing human participants.

**Figure C9.**
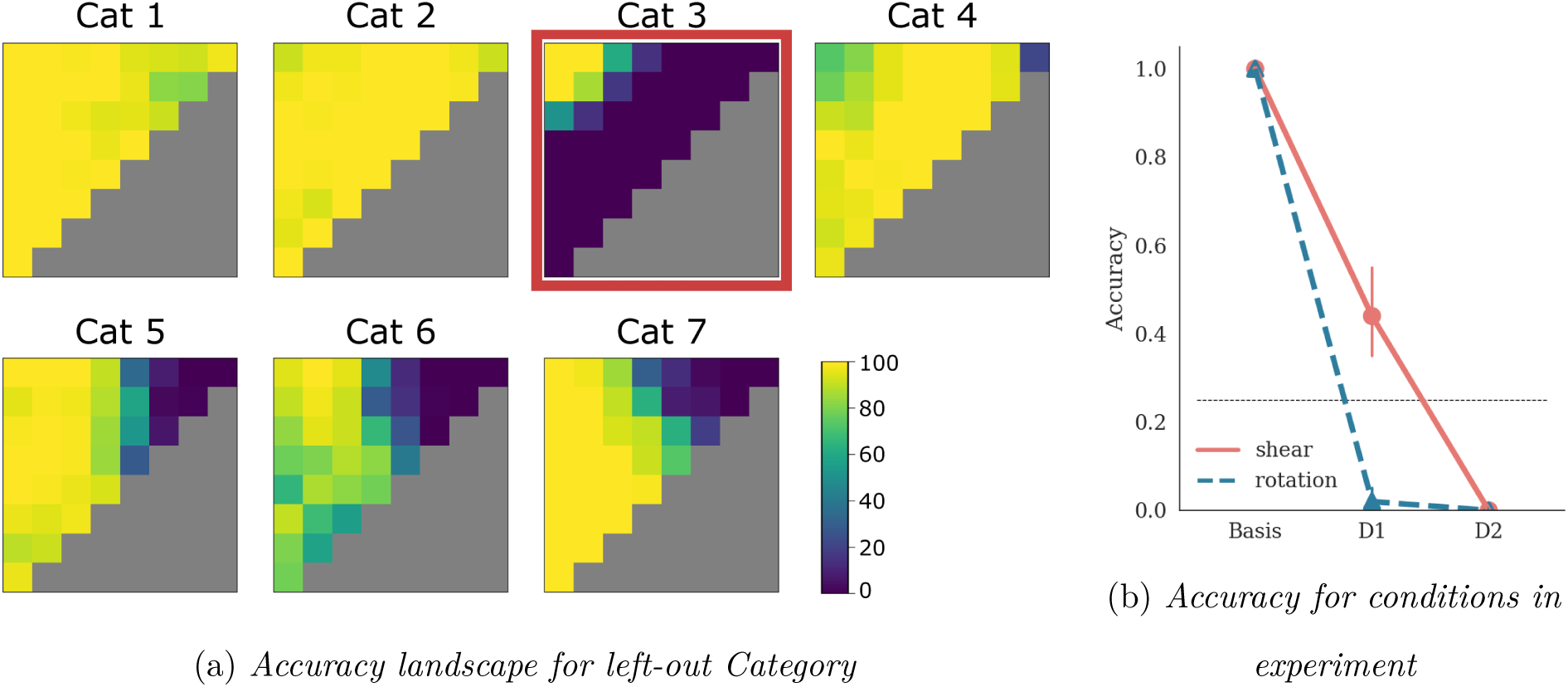
Performance of AlexNet in Experiment 5, Simulation 2. *Note.* Performance of `AlexNet` fine-tuned on an augmented dataset where the basis shapes are not only randomly translated and scaled but also rotated. For six out of seven categories, the network is trained on *all* rotations ([0, 360°)). We then tested the network on the left-out category (Cat 3, highlighted with red square in (a)) on untrained rotations and shears. However, we observed that despite being trained in this manner, the accuracy degraded as a function of the coordinate deformation, rather than the relation deformation. (b) shows the performance of this network for deformations D1 and D2 used to test human participants.

## Footnotes

1 We obtained qualitatively similar results when deformations were organised based on their Euclidean distance to the Basis shape.

2 We did not elicit gender, sex or age information from participants during the experiment and no participant was excluded based on their gender. The proportion of male and female participants reported here is based on the demographics information collected by Prolific when participants register on the platform.

## Notes

This research was supported by the European Research Council Grant Generalization in Mind and Machine, ID number 741134.

### Competing Interest Statement

The authors have declared no competing interest.

### Summary of Updates

We have revised the Abstract and Title for clarity. We have also restructured the experiments and simulations so the Methods of each Experiment / Simulations is reported with the experiment.

